# From expert opinion to empirical evidence: Data-driven parametrization of urban connectivity models using movement-proxy data

**DOI:** 10.1101/2023.12.22.571399

**Authors:** Lisa Merkens, Soyeon Bae, Elin Eberl, Elio Lauppe, Florian Mesarek, Daniel Stauffer-Bescher, Thomas E. Hauck, Wolfgang W. Weisser, Anne Mimet

## Abstract

The landscape connectivity of cities is increasingly recognized as crucial for biodiversity conservation and ecosystem services. Yet, modelling ecological connectivity in cities remains challenging because landscape resistance is often based on expert judgment rather than empirical evidence, leading to varying modelling results and limited use for planning. We developed and tested a data-driven framework for empirically parametrizing resistance and movement-distance parameters in functional connectivity models from movement-proxy data – information on the presence/absence of animal movement from direct observation or camera traps. At each step, we ensured that the connectivity model reflected the behavioural and spatial properties of the observations. We applied the framework for the common blackbird (*Turdus merula*) in Munich, Germany. We used observations of flying blackbirds as movement-proxy data in a logistic regression framework, testing alternative combinations of resistance and movement distances. Model selection identified the parameter sets best supported by the data. The resulting parameters were validated using repeated out-of-sample validation and compared against an expert-based connectivity model. Connectivity derived from empirically estimated parameters increased the probability of observing flying blackbirds. Across repeated validations, the empirical model achieved a mean AUC of 0.76 and R^2^ of 0.17. It performed moderately better than the expert-based model. Depending on their height, buildings exhibited varying resistance to flying blackbirds. Results indicate that expert assessments may oversimplify urban barriers. The approach provides a transparent, reproducible framework for using movement-proxy data to derive maps of landscape resistance. It offers a step toward more data-driven urban connectivity modelling.

## 1. Introduction

Urban ecosystems increasingly serve as important refuges for biodiversity and providers of ecosystem services to people (Kowarik et al. 2025). Green and blue spaces such as parks, gardens, rivers, and remnant woodlands contribute to ecological functioning, mitigate urban heat, and improve human wellbeing (Marselle et al. 2021; Croeser et al. 2021) while preserving biodiversity, including rare and endangered species (Ives et al. 2016; Lepczyk et al. 2023). Recognizing this, international policy frameworks, most notably the Kunming– Montreal Global Biodiversity Framework (Convention on Biological Diversity), explicitly call for increasing the connectivity of urban green spaces to conserve and improve their ability to host biodiversity or provide ecosystem services. Yet, despite growing political attention, cities still lack robust tools to assess and monitor ecological connectivity in ways that meaningfully reflect how wildlife moves through urban landscapes (Habrich and Fahrig 2025).

Ecological connectivity, defined as the degree to which the landscape facilitates or impedes movement among resource patches (Taylor et al. 1993) is central in urban ecology (Goddard et al. 2010). In cities, where habitat is highly fragmented and interspersed with impermeable built structures, maintaining functional connections among greenspaces is crucial for population persistence, gene flow, and daily movements of wildlife (LaPoint et al. 2015). Many species depend on the ability to move across heterogeneous landscapes to forage, breed, or disperse (Goddard et al. 2010; Braaker et al. 2017). Connectivity is also crucial for the delivery of key urban ecosystem services, such as human mental well-being or pollination (Lynch 2019). Thus, understanding and modelling connectivity in urban contexts has become essential for designing ecologically resilient cities.

Over the past two decades, urban connectivity modelling has become a major field within urban ecology and landscape planning (LaPoint et al. 2015; Lookingbill et al. 2022; Habrich and Fahrig 2025). As summarized by Habrich & Fahrig (2025), assessments range from simple metrics of habitat amount to more complex approaches such as functional connectivity and organism-based permeability estimation. Among these, functional connectivity models typically based on least-cost paths or circuit theory have become dominant in urban research because they balance data requirements with biological realism (Calabrese and Fagan 2004). These models typically use multi-class land-cover maps and assign resistance scores to different surfaces based on their assumed difficulty for species to traverse. In urban environments, this framework captures the mosaic nature of the landscape more realistically than binary patch-matrix models (LaPoint et al. 2015; Habrich and Fahrig 2025).

However, a recent systematic review by Habrich & Fahrig (2025) pointed out that many urban connectivity models still assume, rather than demonstrate, that their parameters reflect functional movement. A central challenge arises from the subjective assignment of resistance values (Unnithan Kumar et al. 2022). Resistance scores are commonly derived from expert judgment or literature synthesis (Zeller et al. 2012; Unnithan Kumar et al. 2022; RiordanLShort et al. 2023), which may not accurately capture the true permeability of urban surfaces for a given species. For example, for songbirds in urban environments, maximum expert resistance values range from 50 to 1000 and the relative resistance of streets, buildings and water surfaces varies between studies (Ersoy et al. 2019; Balbi et al. 2021; Bourgeois et al. 2025). This can lead to misleading connectivity maps (Sawyer et al. 2011; Reed et al. 2017) and limits confidence in their application for planning (Habrich & Fahrig, 2025).

At the same time, direct movement data such as GPS tracking, telemetry, or landscape genetics are often unavailable for urban species. These datasets are costly, logistically challenging, and may not align with the temporal or spatial scale of interest (Mateo-Sánchez et al. 2015; Reed et al. 2017). For example, genetic connectivity reflects long-term dispersal and cannot inform planning for short-term movements (Mateo-Sánchez et al. 2015), whereas GPS data can include prolonged resting phases that obscure directed movement (LaPoint et al. 2013; Péron 2019). Therefore, recent studies have explored opportunistic or movement-proxy data, such as roadkill records, capture-recapture data, camera-trap detections, or direct visual or acoustic movement observations (LaPoint et al. 2013; Shimazaki et al. 2016; Mimet et al. 2020; Balbi et al. 2021; Gelmi□Candusso et al. 2025). While Mimet et al. (2020) and Shimazaki et al. (2016) have taken the first steps in demonstrating how these datasets can be used to derive model parameters, they are mostly reserved for post hoc validation (LaPoint et al. 2013; Balbi et al. 2021; Gelmi□Candusso et al. 2025).

Integrating such movement-proxy data directly into model calibration offers a promising way forward. Verbeylen et al. (2003) demonstrated the use of statistical model selection to test multiple resistance scenarios and empirically identify those that best supported by data, but did not use data that reflects actual movement. Using movement-proxy data could enable the derivation of parameters that align with observed movement patterns and can flexibly incorporate diverse data types. However, as emphasized by Habrich and Fahrig (2025), ensuring that the biological process represented by the data matches the spatial and temporal scale of the model is crucial for correct modelling results.

In this study, we extend the approach of Verbeylen et al. (2003) to develop an empirically parameterized functional connectivity model for the common blackbird (*Turdus merula*) in Munich, Germany. Using direct movement observations from urban field surveys as movement-proxy data, we empirically estimate resistance and movement-distance parameters through model selection. We analyse the spatial properties of these movements and adapt the connectivity model to reflect them accurately. Because high-resolution tracking data were unavailable for this species and region, we compared the empirically parameterized model to an existing expert-based model (Ersoy et al. 2019). This comparison allows us to assess whether empirical calibration improves ecological realism relative to the most widely applied approach of expert parametrization. Finally, we discuss how incorporating movement-proxy data into model parameterization can enhance the ecological realism and practical relevance of functional connectivity modelling for biodiversity-informed urban planning.

## 2. Methods

### 2.1 Methods overview

When using empirical data to parameterize connectivity models, it is essential that both the spatial scale (e.g. home-range versus dispersal movements) and the modelled metrics (e.g. patch-versus path-based connectivity) correspond to the ecological processes represented by the data. In this study, we demonstrate how movement-proxy observations, presence or absence of moving individuals, can inform model parameterization when tracking data are unavailable. While detailing the use of model selection to identify the parameters best explaining the data, we also describe how we adapted the connectivity model to ensure that the spatial scales and modelled metrics fit the observation data used for parameterization.

Our workflow comprised six sequential steps, each linking a general modelling decision to a specific implementation in our study (Figure 1). (1) We identify the ecological processes represented by the observation data and the spatial scales at which they operate. (2) Based on this, we define suitable modelling resolution and movement extent. (3) We map resource locations representing potential destinations of movement. (4) We create alternative scenarios for landscape resistance and movement distance. (5) We build a graph-based connectivity model linking resource cells through least-cost paths calculated from the resistance scenarios and prune implausible links according to the distance scenarios. (6) We apply model selection to determine the combination of resistance and distance parameters that best explain the observed movement data. Finally, we compare the empirically calibrated model with an existing expert-based resistance model to evaluate improvements in predictive performance and ecological realism and perform several sensitivity analyses to identify the less robust steps in the modelling workflow. To assess model generality and robustness, we implemented repeated out-of-sample cross-validation, where the parameterization and validation were conducted on randomized subsets of the data.

**Fig. 1.**
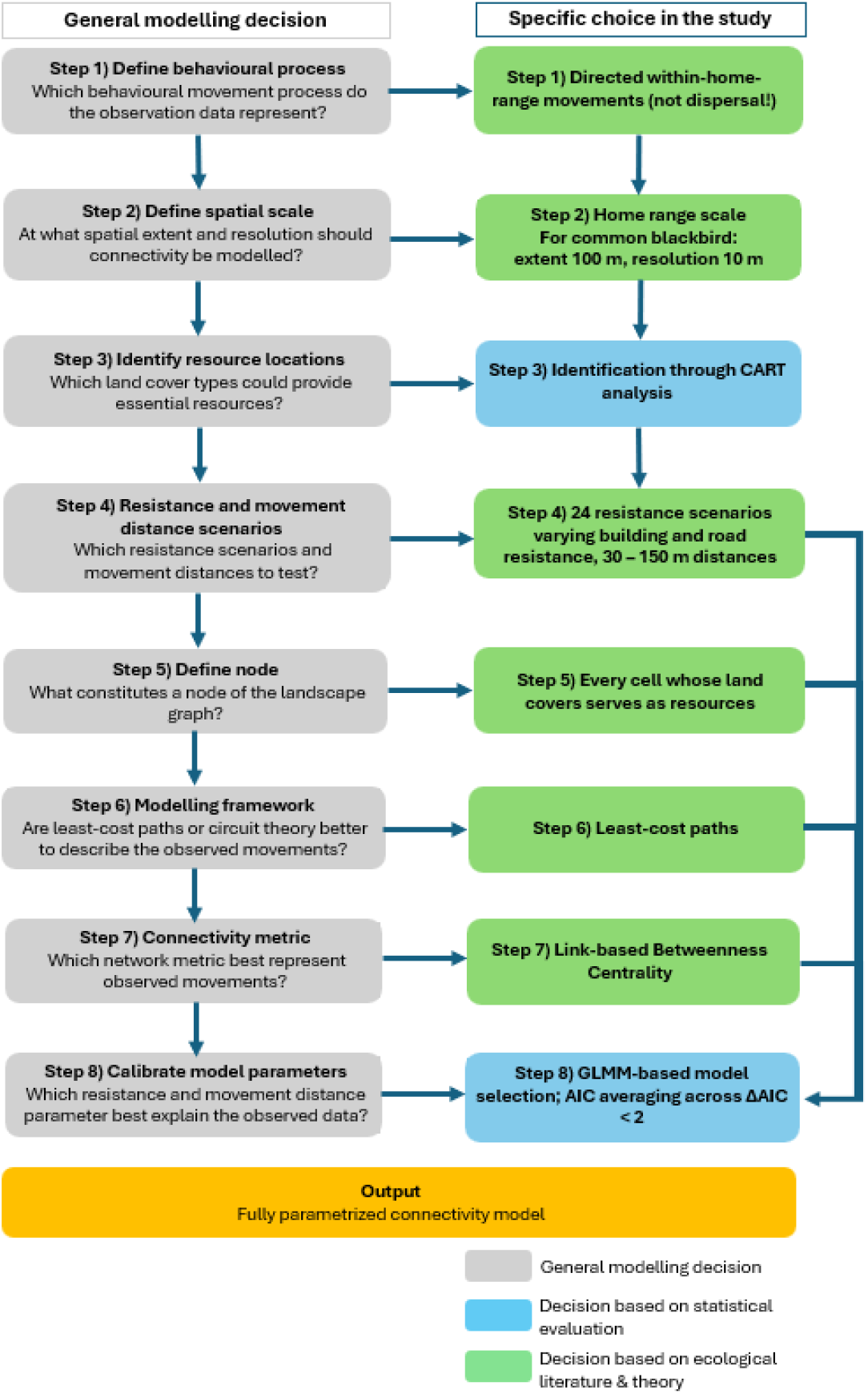
Conceptual workflow for empirical parameterization of a connectivity model and its adaptation to the home range connectivity of the blackbird. Each step (1–6) corresponds to a section of the Methods. The diagram illustrates how general modelling decisions (grey) were implemented through a combination of ecological reasoning (green) and statistical evaluation (blue). Each step ensures that the behavioural and spatial properties of the observation data (i.e. directed within-home-range movements) are reflected in the model structure and parameters. The workflow progresses from identifying the relevant movement process (Step 1) and spatial scale (Step 2), to mapping resources (Step 3), testing resistance and distance scenarios (Step 4), selecting the appropriate network metric (Step 5), and empirically determining resistance and distance parameters via GLMM-based model selection (Step 6). The final output is a fully parameterized, data-driven connectivity model aligned with the ecological processes represented by the observations.

### 2.2 Data sources

#### 2.2.1 Study area and species

Munich (48° 8’ 23” N, 11° 34’ 28” E, 529 m asl) is Germany’s third-largest city with a population exceeding 1.5 million and the highest population density among German municipalities (5100 persons/km2 in 2022) (Stadtverwaltung, n.d.). Despite its urban character, Munich’s green spaces cover approximately 13.4% of the city’s total area with the English Garden encompassing 375 ha that connect the city center with outer areas (muenchen.de, n.d.).

The common blackbird *(Turdus merula)*, an urban dweller frequently found in European cities, was initially a forest species (Mohring et al. 2021). Blackbirds primarily feed on earthworms, invertebrates, and fruits on the ground, in open areas or under canopies (Snow 1966, p. 19). Both males and females establish territories, with suitable nest-sites in trees or shrubs being crucial for territory establishment (Snow 1966). The radius of home ranges for male blackbirds in cities is up to 180 m, but it varies with sex and building density (Snow 1966; Ferry et al. 1981). Due to its distinctive look and song, the common blackbird is an easily identifiable urban species and, therefore, an ideal study subject.

#### 2.2.2 Movement-proxy and species occurrence data

We derived two distinct datasets on *T. merula* occurrence and movement to parameterize and validate the connectivity model (Supplementary Information S1). The datasets differ in behavioural interpretation and spatial sampling design (Table 1).

**Table 1:**
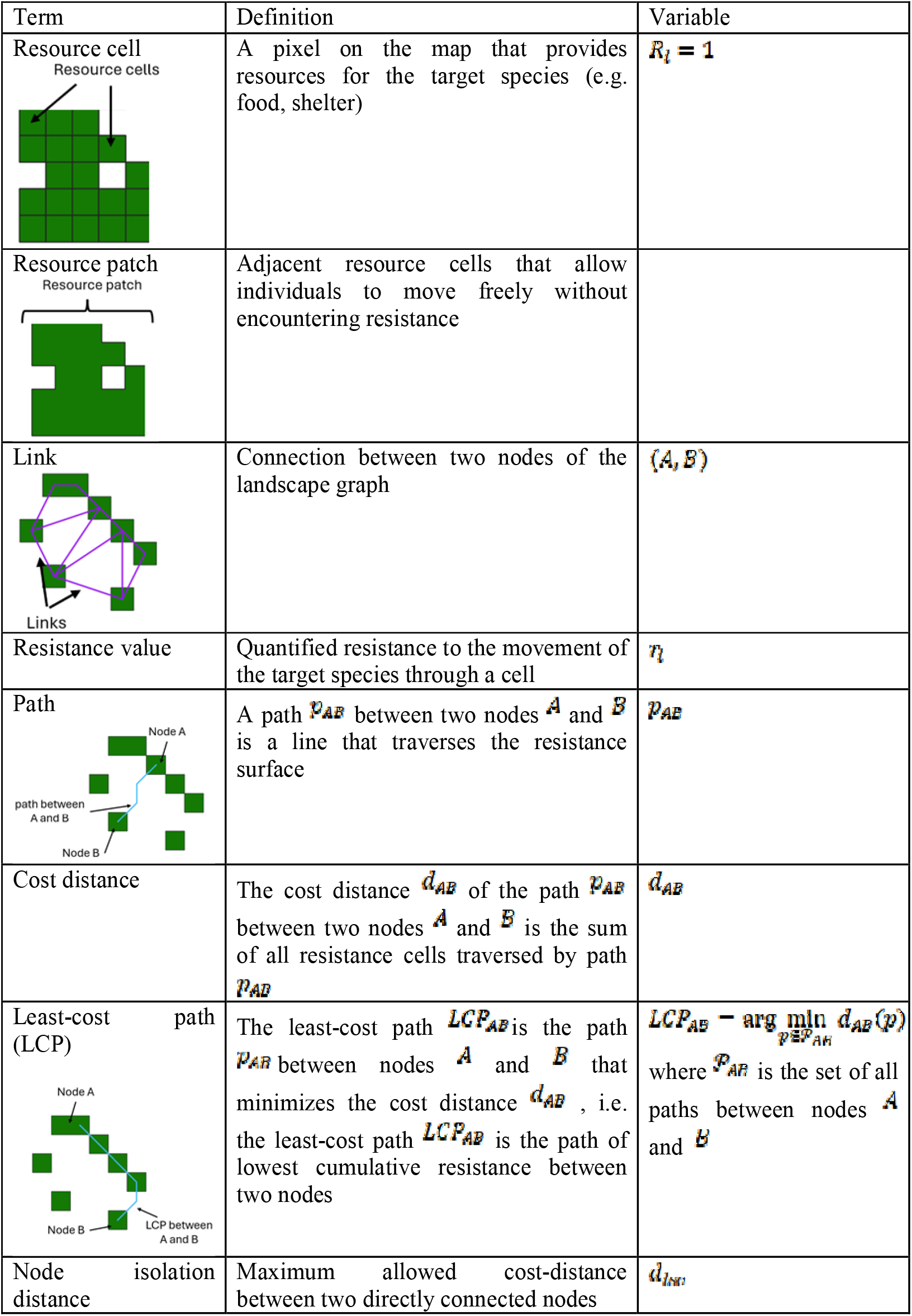
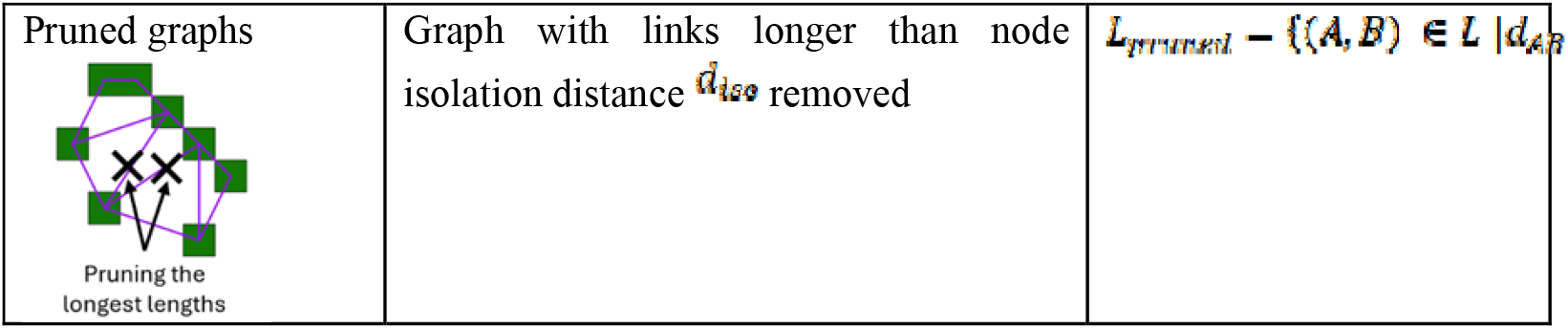
Definitions of the most important terms for model description and their variables.

**Table 2:**
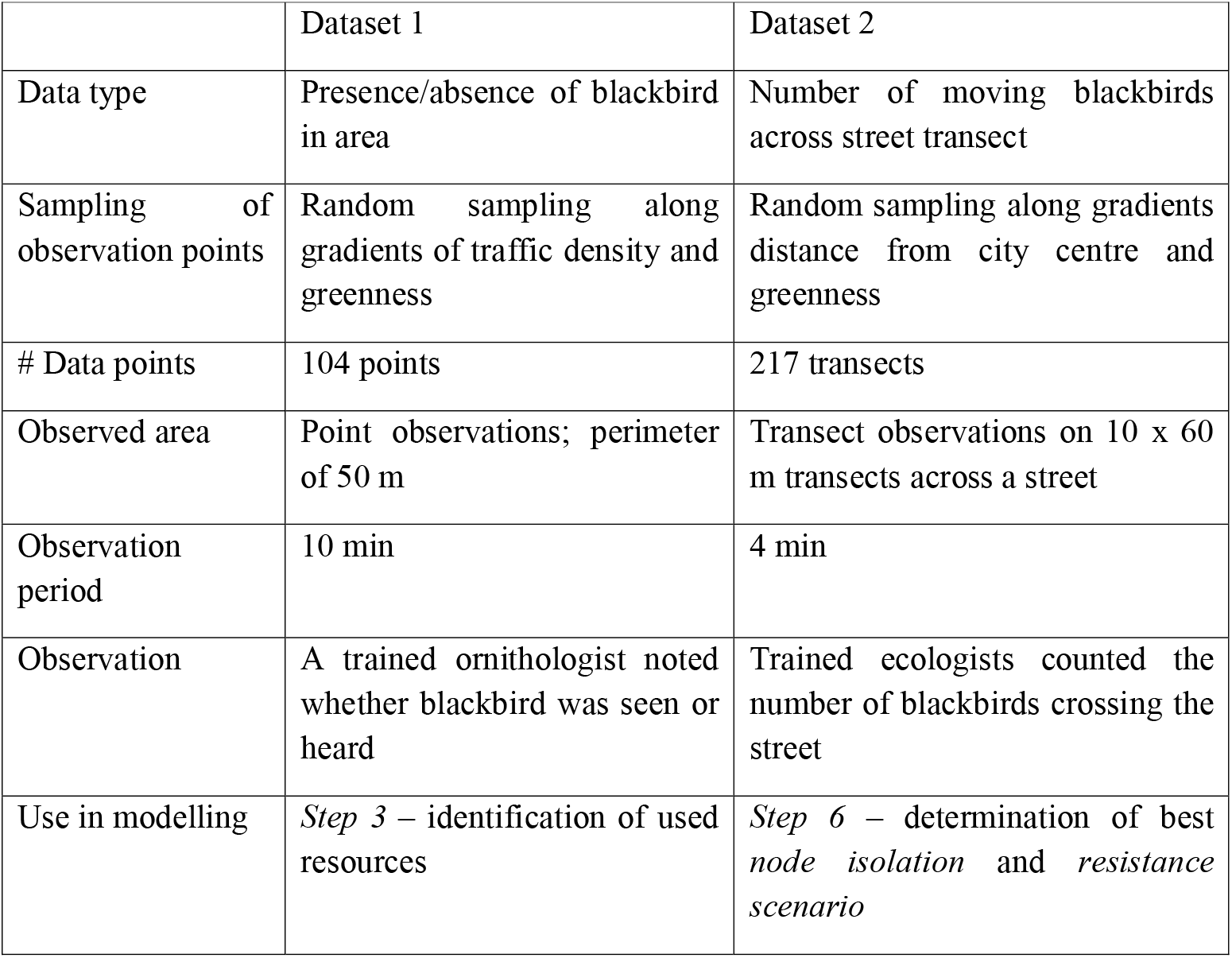
Overview of the datasets used in the parametrization and testing of the connectivity model.

|  | Dataset 1 | Dataset 2 |
| --- | --- | --- |
| Data type | Presence/absence of blackbird in area | Number of moving blackbirds across street transect |
| Sampling of observation points | Random sampling along gradients of traffic density and greenness | Random sampling along gradients distance from city centre and greenness |
| # Data points | 104 points | 217 transects |
| Observed area | Point observations; perimeter of 50 m | Transect observations on 10 x 60 m transects across a street |
| Observation period | 10 min | 4 min |
| Observation | A trained ornithologist noted whether blackbird was seen or heard | Trained ecologists counted the number of blackbirds crossing the street |
| Use in modelling | *Step 3* – identification of used resources | *Step 6* – determination of best *node isolation* and *resistance scenario* |

Dataset 1 comprises visual and acoustic detections representing general presence or absence of blackbirds across the study area. These data were collected through random point observations along gradients of traffic density and greenness and were used in Step 3 to identify resource cells contributing to blackbird occurrence.

Dataset 2 includes observations of blackbirds flying across streets, i.e. locations offering no foraging or nesting opportunities. These flight events were recorded along 10 × 60 m transects and interpreted as directed movements between resource patches. Resources could be nesting, roosting or foraging sites. The dataset was used in Step 6 to estimate resistance values and typical movement distances via model selection. Also, this dataset was used in an out-of-sample cross-validation to compare the performance of our empirically-driven parametrization approach to an expert-driven approach.

#### 2.2.3 Land cover data

We obtained the following high-resolution remote sensing data from the Bavarian State Office for Digitization, Broadband and Surveying (https://www.ldbv.bayern.de/index.html): a digital elevation model at 1 m resolution, a surface model at 40 cm resolution, and color-infrared orthophotos at 20 cm resolution. These were used to create a 40 cm resolution land cover map for Munich in 2017. The map encompasses vegetation classes (grass, shrubs, trees), building height classes (< 10 m, 10 – 18 m, > 18 m), streets, agricultural areas, and water bodies (Supplementary Information S2). These land cover classes are the classes typically considered to be most relevant for urban ecological connectivity modelling (Pundsack et al. 2025).

### 2.3 Step 1) Define behavioural process

Data in Dataset 2 represent home-range movements, rather than long-distance dispersal. Specifically, they capture directed flights between distinct resource patches located within an individual’s typical home-range extent. In this study, a *resource patch* is defined as a continuously vegetated area that allows blackbirds to move freely under canopy or shrub cover without encountering resistance from buildings or vehicular traffic.

These movements between resource patches correspond to the “transit” or “transfer” phase of daily activity, connecting feeding, roosting, and nesting areas (Morales et al. 2004). Because our observations only recorded individuals flying across streets between resource patches, movements occurring entirely within a patch (e.g., short foraging or exploratory movements) (Bailey et al. 1996) were not observed and therefore not modelled. These considerations guided choices for movement scales, modelling resolution, and the choice of graph-based metrics, all tailored to represent functional connectivity of resource patches within the home range rather than dispersal connectivity at the population scale.

### 2.4 Step 2) Define spatial scale

#### 2.4.1 Home range area

Estimates of common blackbird (*Turdus merula*) home range size in urban environments vary between 80 m and 180 m in radius (Ferry et al. 1981; Török and Ludvig 1988; Snow 2008). To represent the most intensively used core area of the home range, we set the home-range radius to 100 m, which defined the search radius for identifying relevant resources in Step 3).

#### 2.4.2 Maximum movement distance in connectivity model

To represent potential commuting movements within this home-range extent, we decided to test potential movement distances of up to 150 m in the landscape graph (Step 4). This upper limit allows for movements crossing substantial parts of the home range, while excluding rare exploratory or dispersal events.

#### 2.4.3 Spatial resolution of connectivity model

To balance ecological realism and computational efficiency, the original 0.4 m land-cover data were aggregated to 10 m cells, calculating the proportion of each land-cover class per cell. This resolution captures the fine-scale heterogeneity relevant to blackbird foraging and movement decisions (e.g., shrubs, small lawns) while reflecting the species’ perceptual and locomotor scale (Supplementary Information S3). At this resolution, several resource cells fit within a single home range, maintaining ecological detail while keeping model complexity manageable.

### 2.5 Step 3) Identify resource locations

The first step in constructing the connectivity model was to identify which land-cover types constitute relevant resources for the common blackbird (*Turdus merula*) and to map their spatial distribution across the study area. This mapping defined the potential destinations of movement and determined which cells could later serve as nodes in the landscape graph.

To identify resource types associated with blackbird occurrence, we used Dataset 1, which contains presence/absence observations across urban gradients in Munich. We hypothesized that specific vegetation features, such as trees, grass and shrubs, increase the probability of blackbird presence, whereas impervious surfaces would have weaker or negative effects. For each observation point, we calculated the proportion of tree, shrub, and grass cover within a 100 m radius, corresponding to the estimated home-range radius (Section 2.4).

We applied Classification and Regression Trees (CARTs) to model blackbird presence as a function of these potential resource types. CARTs were chosen for their ability to handle categorical response variables, non-linear relationships, and interactions among predictors, while maintaining transparent and interpretable results (De’ath and Fabricius 2000; Bel et al. 2009). In contrast to more complex ensemble methods such as Random Forests, CARTs allow clear ecological interpretation of threshold responses and are easily translated into landscape classifications (Cutler et al. 2007; Pesch et al. 2011; Mimet et al. 2019). Model performance was evaluated using a confusion matrix based on the training data.

Applying the fitted CART model to the complete study area identified areas where vegetation cover exceeded the minimum resource requirements for blackbird occurrence. Given the generally high vegetation cover in Munich, nearly all areas met these thresholds. We therefore retained the entire city for connectivity modelling.

Within these suitable areas, we designated resource cells as all 10 m pixels containing at least 20 m^2^ of the land cover identified as a resource according to the CART, following the considerations of Bourgeois et al. (2024). For example, if the CART identified shrubs as an important predictor for blackbird occurrence, all cells with a minimum shrub cover of 20 m^2^ became graph nodes. The resulting resource map delineated the spatial distribution of resource cells (R□ = 1) and non-resource cells (R□ = 0) and formed the foundation for constructing the landscape graph described in Section 2.7.

### 2.6 Step 4) Resistance and movement-distance scenarios

Resistance values in connectivity models summarize multiple behavioural processes: movement preferences, avoidance, energetic costs, perceptual limits, and mortality risk (Zeller et al. 2012). We adopted a scenario-based parameterization combined with model selection (Verbeylen et al. 2003). This approach tests multiple ecologically plausible combinations of resistance and movement-distance parameters and identifies those that best explain observed movement patterns (Dataset 2).

#### 2.6.1 Resistance scenarios

We generated a set of candidate resistance maps ℛ_1_, …, ℛ_*n*_ by varying the resistance values assigned to different land-cover types. In each scenario, resource cells identified in Section 2.5 were assigned a resistance value of 1, representing undisturbed movement. Vegetation not classified as a resource, as well as agricultural and open water areas, were assigned a moderate resistance of 10 to reflect their open and less structured nature (Tremblay and St. Clair 2009). Because Dataset 2 contained few observation points near water bodies or agricultural land, their influence on movement could not be estimated directly from the data; therefore, their resistance values were defined a priori.

Anthropogenic structures such as streets and buildings were assumed to impose stronger barriers due to increased predation risk, energetic cost, and reduced permeability (Tremblay and St. Clair 2009, 2011; Shimazaki et al. 2016). We therefore assigned them resistance values higher than those for vegetation, ranging from 10 to 1000 (Bourgeois et al. 2024). To represent ecological realism, taller buildings were always assigned equal or greater resistance than lower buildings, but no prior assumption was made about their resistance relative to roads.

This factorial design yielded 24 distinct resistance scenarios, encompassing all combinations of plausible relative resistance orders under these constraints. Each resistance map was produced by summing the weighted contributions of land-cover types within each 10 m cell of the land-cover map, yielding a continuous resistance surface consistent with the spatial resolution defined in Section 2.4.

#### 2.6.2 Movement-distance scenarios

We further defined a set of candidate node-isolation distances *D* = {30, 60,90, 150m}, representing the maximum distance over which direct movement between two resource cells is assumed possible. These distances were selected based on reported within-home-range movement distances and home-range radii of urban blackbirds (Ferry et al. 1981; Török and Ludvig 1988; Cresswell 1999; Snow 2008).

Combined, these 24 resistance and four distance scenarios yielded 96 resistance-distance scenarios tested through model selection (Section 2.8).

### 2.7 Graph-based connectivity model

#### 2.7.1 Step 5) Define node

To represent fine-scale functional connectivity within blackbird home ranges, we implemented a cell-based landscape graph rather than a traditional patch-based approach. In patch-based graphs, multiple contiguous resource cells are aggregated into single habitat patches, and movement is assumed to occur only between their edges. By contrast, in a cell-based graph, each resource cell acts as an individual node, and movements can occur both within and between resource clusters (Table 1) (Figure S3). This finer spatial representation allows for a more continuous depiction of potential movement pathways (Drielsma et al. 2007; Saura and Pascual-Hortal 2007).

Cell-based graphs are particularly suitable for representing within-home-range movements, such as the directed flights observed in our data (Dataset 2). These movements typically connect nearby resource sites, trees, shrubs, or small vegetation patches, across short distances (tens to hundreds of metres) and through partially permeable barriers like streets or courtyards. Aggregating these features into large patches would obscure the fine-scale resource heterogeneity and redundant paths between patches available to individuals. As supported in recent studies (Goicolea et al. 2021; Van Moorter et al. 2023), such redundant pathways are a critical feature of realistic connectivity models, as they reflect the diversity of movement choices animals can make within complex landscapes.

Nodes in our graph corresponded to resource cells identified in Section 2.5. The 10 m grid resolution (Section 2.4) thus defined both the spatial extent and size of individual nodes.

#### 2.7.2 Step 7) Modelling framework

To represent potential movement routes between resource cells, we applied the least-cost path

(LCP) framework (Adriaensen et al. 2003). LCPs estimate the route between two locations that minimizes the cumulative resistance, defined as the total “cost distance” *d*_*AB*_ across all traversed cells (McClure et al. 2016). This approach assumes that animals move through their familiar environment in a manner that minimizes energetic expenditure and exposure to risk, selecting paths of lowest cumulative cost (McClure et al. 2016). In our case, this approach is particularly appropriate because long, directed home-range movements, such as those observed for blackbirds moving between resource sites within their territory, occur within a well-known spatial context. Individuals are likely to rely on spatial memory to optimize travel routes, avoiding energetically expensive or dangerous areas such as roads and built structures (McClure et al. 2016; Blazquez-Cabrera et al. 2016). Unlike circuit-theory approaches, which simulate undirected random walks and are better suited to dispersal-scale movement (McRae 2006), LCPs explicitly represent directed movements that link known resource locations.

In graph-based connectivity models, including all possible links between nodes can result in unrealistic or biologically implausible movement pathways, particularly when modelling movements that occur within an animal’s home range. Animals typically move between nearby resource sites or “stepping stones” and avoid long, exposed crossings without cover (Bélisle and Desrochers 2002). Consequently, unrestrained link generation may create paths that greatly exceed the spatial extent of realistic movements, thereby inflating connectivity estimates and computational demands (Mimet et al. 2016). Following established practice in landscape graph analyses (Foltête et al. 2012b), all links with a cumulative cost distance *d*_*AB*_ > *d*_iso_ were removed from the graph.

From each set of least-cost paths generated under the 24 resistance scenarios (Section 2.7.2), we constructed four pruned graphs. Each graph corresponded to one of the four candidate movement distances: 30, 60, 90, and 150 m, reflecting the range of empirically plausible within-home-range movement distances for urban blackbirds.

#### 2.7.3 Step 7) Connectivity metric

We computed the link-based Betweenness Centrality (BC) that identifies the edges most central to the network’s overall flow, i.e. those expected to be traversed most often under the least-cost assumptions (Girardet et al. 2015). Because our empirical data capture movements between resource patches (Dataset 2) rather than occupancy or activity within patches, link-based metrics are most appropriate for reflecting the ecological process under study (Girardet et al. 2015). If observations were limited to presence or absence within patches, node-based metrics could more directly represent connectivity. Formally, for a given link *l*, BC is defined as

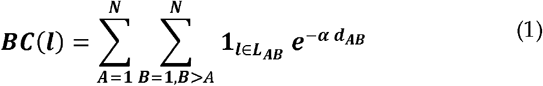

Where *L*_*AB*_ is the set of links traversed by the least-cost path between nodes *A* and *B* and the indicator function 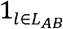 ensures that only links occurring on the path are included. The weighting term 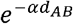 reflects movement probability along the path with cost distance *d*_*AB*_, with the decay parameter *α* defined as:

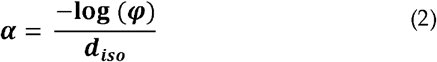

so that movement probability drops to *φ* at node isolation *d*_*iso*_ (Bodin and Saura 2010). Typical values for *φ* are 0.05 or 0.01 (Bodin and Saura 2010).

The graph-based model was constructed using the Graphab 2.8 software (Foltête et al. 2012a) via the graph4lg R package v. 1.8 (Savary et al. 2023). For each combination of resistance and movement distance, we generated least-cost graphs and calculated link-based BC values. To parameterize the connectivity model using statistical model selection, it was necessary to obtain continuous connectivity values for all parts of the study area. (Foltête et al. 2012b). Therefore, we interpolated the BC values from links by adapting Graphab’s interpolation workflow to a continuous surface representing the entire landscape.

### 2.8 Step 8) Calibrate model parameters

#### 2.8.1 Model selection

To empirically determine the most plausible resistance values and movement distances, we applied a model selection procedure comparing the predictive performance of connectivity maps derived from different combinations of resistance and node-isolation scenarios (Verbeylen et al. 2003). For each combination of resistance scenario *R*_*k*_ and node isolation distance *d*_*l*_, we constructed a corresponding landscape graph and generated a continuous connectivity map *BC*^(*k,l*)^ (*x,y*)(Section 2.7.5). This parameterization procedure was repeated within each out-of-sample training subset of the validation (section 2.9), ensuring that resistance and movement parameters were estimated independently for every training–test split.

From these maps, we extracted connectivity values *BC*^(*k,l*)^ (*x*_*i*_,*y*_*i*_) at the observation locations where blackbird movements were recorded (Dataset 2). Each observation represented the presence (1) or absence (0) of movement across a street segment. The probability of observing movement at each location was modelled as a function of the predicted connectivity and relevant local covariates using the following generalized linear mixed model (GLMM):

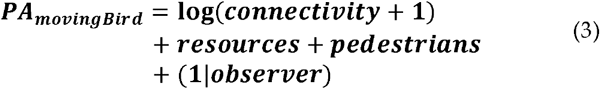

where connectivity is the mean interpolated BC value within the observation transect, resources the proportion of resource cells, and pedestrians the number of pedestrians counted during data collection. We accounted for the number of pedestrians during observation to capture the disturbance that the flying blackbirds could experience, which is not directly covered by the sources of resistance in the connectivity model. The random intercept for observer accounts for potential observer-related variation in detection.

Model fitting was conducted in R using the lme4 package (Bates et al. 2015). We used DHARMa (Hartig 2022) to test for outliers, overdispersion, zero inflation and homoscedasticity.

#### 2.8.2 Best resistance and movement distance values

To identify the combination of resistance and node isolation distance that best predicted observed blackbird movements, we compared all 96 GLMMs (24 resistance scenarios × 4 isolation distances) using the Akaike Information Criterion (AIC) (Verbeylen et al. 2003). The most parsimonious parameter set 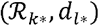 minimized the AIC across all models. Within each repetition of the out-of-sample cross-validation, the best-performing parameter combination was recorded, and resistance and distance values were averaged across the top-ranked models before validation. This procedure allowed us to determine which combination of resistance and distance values provided the best fit to the empirical movement data, thereby directly estimating parameter values rather than validating pre-defined ones.

Because model uncertainty may remain when several parameter combinations perform similarly well, we applied an evidence-based model averaging approach. All models within a ΔAIC ≤ 2 of the best-performing model were considered statistically equivalent with differences in statistical performance being too low to identify one best model (Grilo et al. 2011). To account for model selection uncertainty, we averaged the resistance and node-isolation distance values across these top-ranked models, following the principle of AIC-based model averaging (Anderson and Burnham 2002):

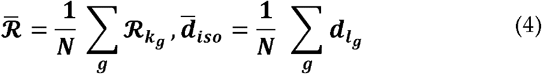

where *g* indexes the models within the ΔAIC ≤ 2 range and *N* is their number. This approach provides parameter estimates that are less sensitive to small variations in data and more robust to the choice of any single best-fitting model (Anderson and Burnham 2002). The resulting averaged resistance map 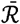 and isolation distance 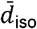 represent the empirically derived parameterization that best explained the observed movement probabilities.

### 2.9 Validation and model comparison

#### 2.9.1 Monte Carlo cross-validation

To evaluate the predictive performance and robustness of the empirically parameterized connectivity model, we implemented a repeated out-of-sample validation (also known as Monte Carlo cross-validation). Prior to the final model validation, we assessed the robustness of the parameterization to varying data sizes by repeating the full procedure with 33 % and 66 % of the total Dataset 2 (Supplementary Information S4). Parameter estimates for resistance and movement distance remained consistent across these reduced-sample analyses, indicating that the calibration approach is stable even with substantially fewer data points. Based on this robustness, we opted for a 50:50 train–test split in the repeated out-of-sample validation. This design provides a larger independent test set, thereby reducing the influence of individual or spatially clustered observations on validation outcomes while maintaining sufficient data for reliable parameter estimation.

The full dataset of movement observations was randomly partitioned ten times into training (50 %) and test (50 %) subsets. In each repetition, the complete parameterization procedure (Sections 2.6–2.8) was applied exclusively to the training data to derive resistance and node-isolation parameters. These parameters were then used to construct a connectivity model and generate an interpolated Betweenness Centrality (BC) map, which was evaluated against the independent test data. This repeated random-subsampling approach provides robust estimates of model performance and parameter stability while avoiding over-fitting to a single partition of the data.

For the withheld test data, mean BC values were extracted at each observation transect, and the probability of observing blackbird movement was predicted using the GLMM already used for parametrization (equation 3). Predictive accuracy was quantified by the area under the receiver-operating-characteristic curve (AUC) using the pROC R package (Robin et al. 2011) and by *R*^2^ from glmmTMB package (Brooks et al. 2017), computed for each repetition (Grilo et al. 2011). The ten repetitions hence provided independent estimates of model performance and variability arising from data partitioning.

#### 2.9.2 Comparison with expert-based model

We further compared our empirically parameterized model with an expert-driven model for the common blackbird published by Ersoy et al. (2019). Based on their expert-derived resistance values (resistance = 1 for trees and shrubs, resistance = 25 for all other land-cover classes), we created a corresponding resistance map for Munich. To isolate the effect of resistance values while keeping all other parameters constant, we repeated our modelling workflow using the expert-derived resistance map of Munich but applied the node-isolation distance identified from our data-driven parameterization within the same repetition. Otherwise, we strictly followed our modelling workflow as described in section 2.7. This comparison directly quantifies the added predictive value of empirically derived resistance values relative to expert-defined ones.

#### 2.9.3 Performance metrics and synthesis

For each repetition, we summarized model performance by reporting AUC and *R*^2^ values for (1) the model with expert resistance and (2) our full empirically-derived model. The resulting distributions across the ten repetitions provided a measure of both the consistency and magnitude of these performance differences. Finally, the resistance values and node-isolation distances estimated across all repetitions were averaged to obtain a final, data-driven parameter set used for generating the overall connectivity map presented in Section 3.

### 2.10 Sensitivity analyses

Sensitivity analyses were conducted to identify parameters that strongly influence the outcome of the connectivity modelling (Cariboni et al., 2007; Rayfield et al., 2010). We evaluated the robustness of the parametrization procedure to changes in the home-range radius used for resource identification and the spatial resolution of the input data (Supplementary Information 6). o test the effect of home-range definition, we repeated the CART analysis using radii of 80 m and 120 m instead of 100 m. To evaluate the influence of spatial resolution, we compared the selection of model parameters obtained at modelling resolutions of 6 m and 20 m.

## 3. Results

### 3.1 Results of data-driven connectivity model

#### 3.1.1 Resource identification

The CART analysis predicting blackbird occurrence from resource cover achieved a prediction accuracy of 0.82 and a Kappa value of 0.57, indicating good model performance beyond random expectation (Donker et al. 1993). Blackbird occurrence increased with tree cover exceeding 16% or shrub cover above 1.7% within a 100 m radius. Grass cover showed no significant relationship with occurrence (Supplementary Information S5). The results were robust to alternative home range radii (80 m, 120 m), with trees and shrubs consistently identified as key predictors of occurrence.

#### 3.1.2 Parameterization of resistance and movement distance

Mean node isolation distances and resistance values were obtained by averaging across the best models within ΔAIC < 2 per repetition. The resulting mean node-isolation distance was 83 m, corresponding to the typical range of long, directed movements within blackbird home ranges. Empirically derived resistance values indicated that high-rise buildings posed the strongest barrier to movement (mean resistance = 427), followed by medium-rise buildings (128). Low buildings had a mildly higher resistance (56) than streets (52). These results suggest that among the urban land cover types, especially high and medium-sized buildings differ in their resistance to blackbird movement relative to low buildings and streets. This relative difference was not accounted for by the expert-based resistance (Ersoy et al. 2019). Additionally, all empirically derived resistances were higher than the expert-based values (25) reported by Ersoy et al. (2019).

The relative order of resistance values was additionally supported by the frequency with which different resistance maps were selected. Model selection frequencies revealed high consistency for tall buildings, where most top models supported very high resistance (1000) and few supported lower values. For medium buildings, intermediate to high resistance (100– 1000) was most frequently selected. Resistance of streets and low buildings exhibited greatest uncertainty, with values of 10, 100, and 1000 all appearing among the best models (Figure 2a). Node-isolation distances were also variable with highest frequency of intermediate movement distances (Figure 2b).

**Fig. 2.**
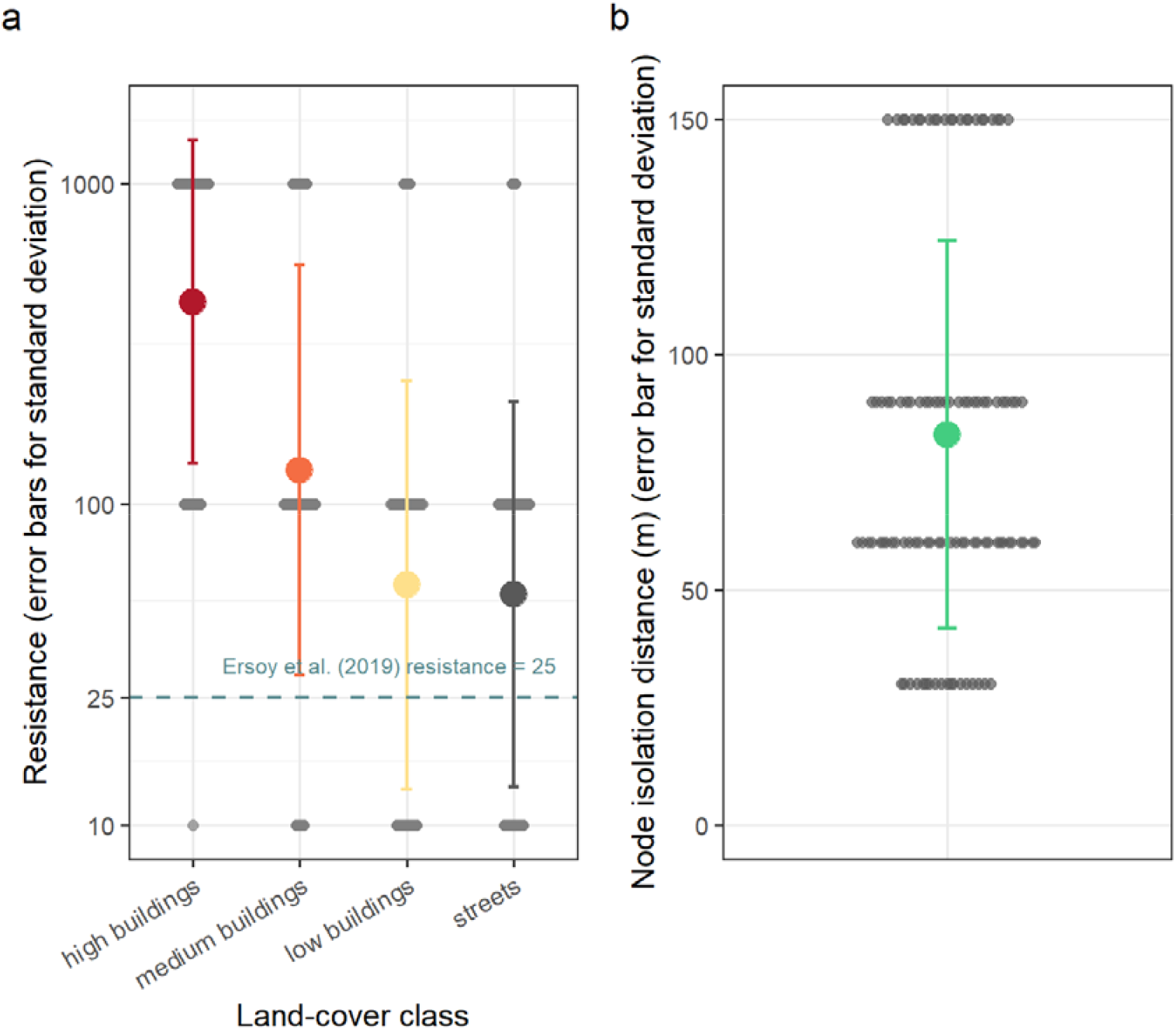
Empirically derived movement parameters for the common blackbird (*Turdus merula*) across ten out-of-sample cross-validation repetitions. (a) Mean resistance values (± SD) for the four urban land-cover classes included in model calibration, displayed on a logarithmic scale. (b) Mean node-isolation distance (± SD) representing the spatial scale of movement most consistent with the observed street-crossing data. Individual points indicate values estimated for each repetition. The dashed line in panel (a) marks the uniform resistance value (25) used in the expert-based model of Ersoy et al. (2019).

#### 3.1.3 Spatial patterns of connectivity

The resulting connectivity map (Figure 3c) showed high connectivity in rural areas and suburban zones with dense shrub and tree coverage. Additionally, connectivity was high around the large green spaces and rivers within the city centre, particularly within the English Garden in Munich’s northeast and along the Isar River in the southeast. However, connectivity was lowest in the dense urban core characterized by tall buildings and impervious surfaces. Agricultural areas at the city fringes also showed low connectivity.

**Fig. 3.**
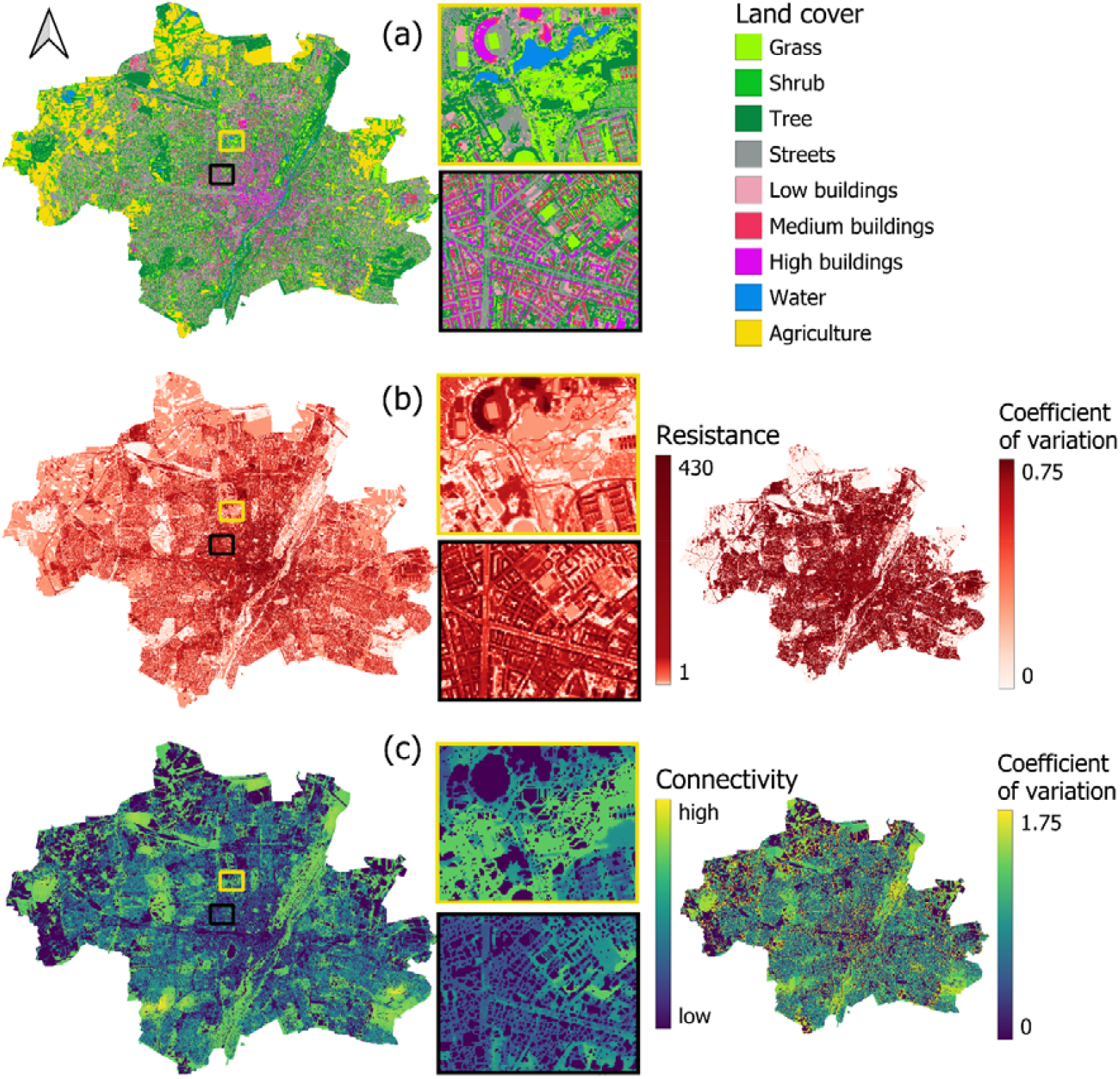
Land cover, empirically derived resistance, and resulting connectivity surfaces for the common blackbird (*Turdus merula*) in Munich, Germany, including associated uncertainty. (a) Land-cover classification used for resistance parameterization and connectivity modelling, with zooms showing suburban (yellow frame) and dense urban (black frame) areas. (b) Mean resistance surface (left) and its coefficient of variation (right) across cross-validated models. (c) Mean connectivity surface (left) and its coefficient of variation (right) across alternative resistance and distance parameterizations. The coefficient of variation (CV) expresses the relative variability among model outputs (CV = SD / mean), providing a normalized measure of uncertainty. High CV values indicate areas where predicted resistance or connectivity is most sensitive to model assumptions, while low values denote stable and robust estimates.

Areas of low uncertainty in connectivity were primarily found where overall connectivity was low, particularly within the dense urban core and the surrounding agricultural landscapes. In contrast, highest uncertainty occurred where connectivity was concentrated along narrow or isolated pathways, such as corridors crossing arable fields. In general, areas of high predicted connectivity also exhibited greater variability among model runs. This mainly indicates that the absolute magnitude of connectivity in these zones may be uncertain. Nevertheless, their relative importance as movement corridors remains robust.

#### 3.1.4 Model performance

Connectivity calculated from empirically derived parameters increased the probability of observing blackbird movements across in nine out of ten validation repetitions (Figure 4).

**Fig. 4.**
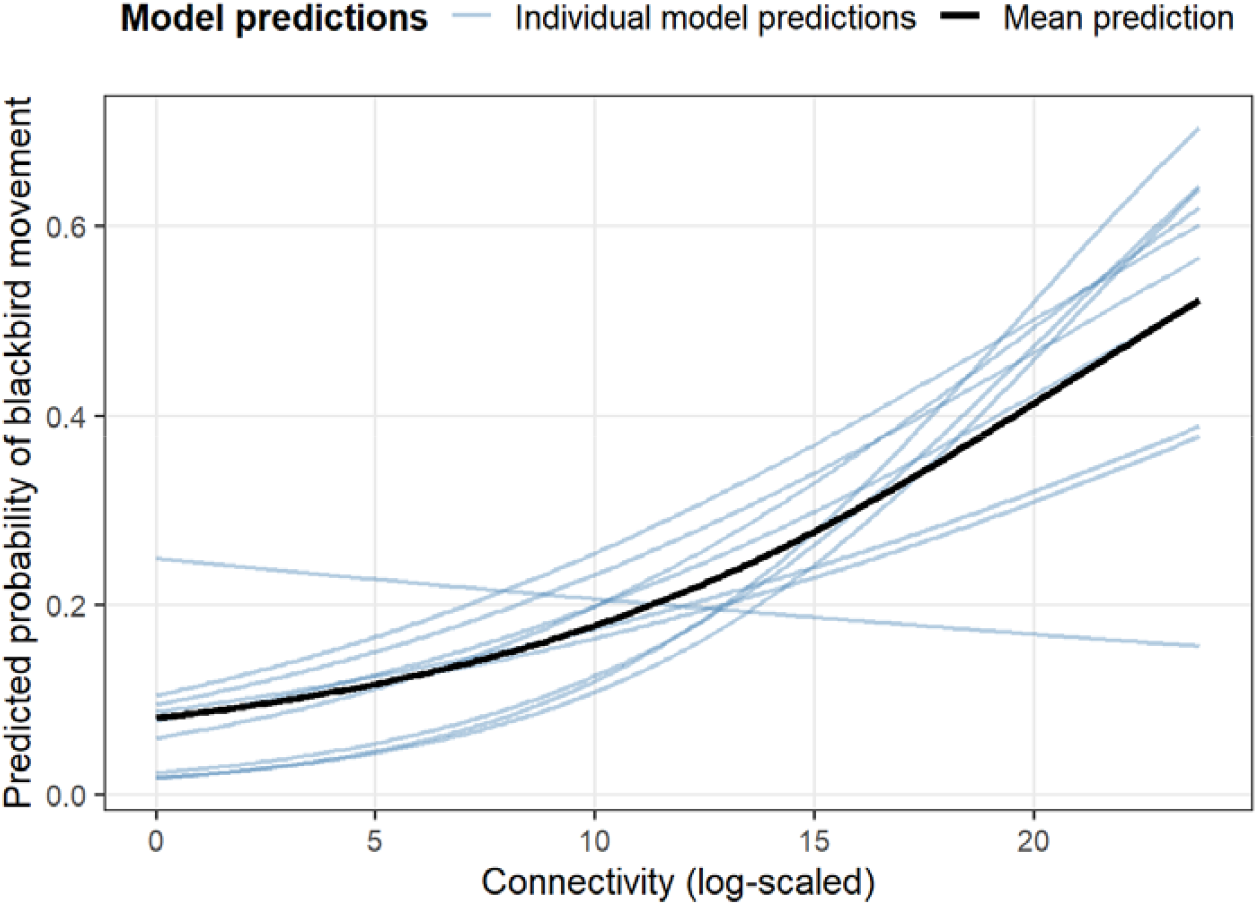
Relationship between empirically derived connectivity and blackbird occurrence probability. Blue lines show predictions from each of the ten out-of-sample repetitions; the black line shows the mean relationship.

The empirical model achieved a mean AUC of 0.76 (range 0.69–0.85) and a mean R^2^ of 0.17 (range 0.02-0.32) across the ten repeated test-training splits, indicating acceptable to good discrimination ability (Hosmer et al. 2013). In nine out of ten GLMMs, the coefficient for log-transformed connectivity was positive, confirming that areas with higher modelled permeability were more likely to contain movement observations. The mean effect size corresponded to roughly a 13% in the odds of observing blackbird movement for each one-unit increase in log(connectivity), highlighting a consistent positive association between modelled connectivity and observed movement patterns. As shown in Figure 4, predicted probabilities rose steadily with increasing connectivity, and while repetition-specific responses varied in steepness, all but one showed a clearly positive trend.

### 3.2 Comparison with expert-based model

The repeated out-of-sample validation revealed that our fully empirical model performed moderately better than the variant using expert resistance (Ersoy et al. 2019) (Figure 5). On average, AUC and R^2^ values were 0.012 higher for the empirical model than for the expert-based model, indicating a small but consistent performance gain. This suggests that the expert-based model also performs acceptably, but that the empirically-based model provides additional information relevant to discriminate between areas of blackbird movement and those without.

**Fig. 5.**
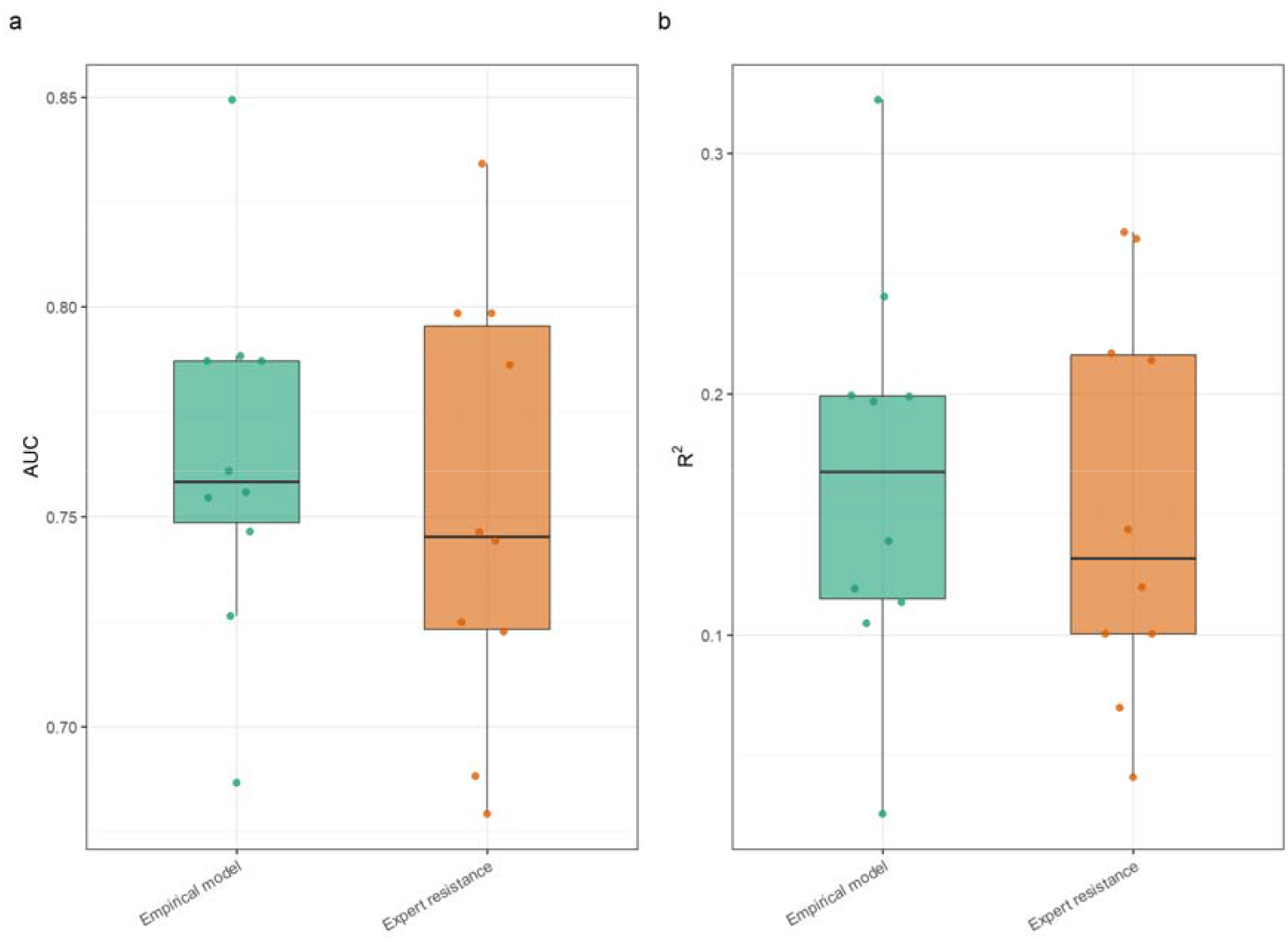
Results of the repeated out-of-sample validation (Monte Carlo cross-validation) comparing empirical and expert-based connectivity models. (a) AUC and (b) R^2^ values for the ten validation repetitions. The empirical model was parameterized using movement-proxy data, while expert-based models used resistance from Ersoy et al. (2019). Boxes indicate interquartile ranges and points represent individual validation repetitions.

## 4. Discussion

Our results demonstrate that empirically calibrating resistance and movement-distance parameters yields connectivity models that are ecologically consistent with observed movement behaviour and modestly outperform expert-based approaches in predictive accuracy. While the overall improvement in AUC and R^2^ was moderate, the empirically parameterized approach provides a stronger process-based justification for the resulting resistance values and movement scales. The consistency of parameter estimates across repetitions further supports the robustness of the derived resistance maps, while the model selection framework allows the spatial delineation of areas with higher or lower confidence in simulated connectivity. Together, these results underscore the importance of data-driven calibration for enhancing the ecological realism, reproducibility, and interpretability of functional connectivity models in urban landscapes.

### 4.1 Ecological realism of empirically derived movement parameters

The empirically derived parameters are consistent with established knowledge of blackbird ecology, suggesting that the calibration procedure captured ecologically meaningful movement processes. The positive effects of shrubs and trees, but not grass, on blackbird occurrence align with previous studies from both rural and urban contexts (Mohring et al. 2021). The estimated node-isolation distance of approximately 80 m corresponds closely to the typical length of directed movements observed in wild blackbirds foraging or commuting between feeding and nesting sites (Török and Ludvig 1988; Cresswell 1999). Together, these findings suggest that the modelled movement scale accurately represents movements within the home range rather than long-distance dispersal.

Urban land-cover types differed markedly in their estimated resistance. High-rise buildings exhibited the strongest resistance to movement, followed by medium-rise and low buildings. This pattern implies that dense, vertical urban structures pose substantial barriers to blackbird movements. The high resistance could reflect increased energetic costs and mortality risks when crossing built-up areas. The results are consistent with previous studies showing that urban birds avoid highly sealed or exposed areas and that buildings inhibit movement more strongly than other sealed surfaces (Tremblay and St. Clair 2009; Balbi et al. 2021). The model selection procedure consistently assigned high resistance to tall and medium buildings, suggesting that these estimates are relatively robust across repetitions. In contrast, resistance values for streets were more variable. This variability could reflect the heterogeneity within the “street” class, which includes a broad range of sealed surfaces differing in traffic intensity and other disturbances. The validation of an urban connectivity model for passerines by Balbi et al. (2021) supports that streets of high traffic density have higher resistance to bird movement. Consequently, the model may have alternated between different resistance values for streets between repetitions, depending on the local characteristics of the training dataset.

Overall, empirically derived resistances were substantially higher than those that experts suggested for the *Turdus merula* in the study by Ersoy et al. (2019). This could indicate that the expert map may have underestimated the difficulty of movement through built environments and oversimplified the resistance values of different urban land covers. This divergence highlights a key advantage of data-driven calibration: even for urban-tolerant species such as the blackbird, expert judgement may yield overly optimistic assumptions about matrix permeability. In contrast, empirically estimated resistance values better reflect the actual behavioural constraints imposed by the urban matrix. The resulting resistance and connectivity maps depict realistic spatial gradients in movement cost, with highest permeability along vegetated suburban corridors, riparian strips, and large urban parks, and lowest permeability in the dense city centre dominated by tall buildings and impervious surfaces.

Model performance metrics further support the ecological plausibility of the empirically parameterized model. The proportion of variance in movement probability explained by connectivity is, for many repetitions, comparable to those reported for other urban species using tracking data (Braaker et al. 2014). This confirms a good predictive capacity given the complexity of behavioural data. Connectivity had a consistently positive effect on movement probability across nine out of ten cross-validation folds, indicating that long, directed blackbird movements between resource patches are shaped by both structural barriers and resource distribution (Ersoy et al. 2019; Balbi et al. 2021). The model therefore captures key ecological processes governing blackbird movement in urban environments and underscores the importance of small-scale connectivity for maintaining functional movement pathways under variable resource availability.

### 4.2 Comparison with expert-based model

Including the expert-based resistance map of Ersoy et al. (2019) within our model selection procedure provided a direct empirical test of its performance compared to the approach selected in many urban connectivity studies (Habrich and Fahring 2025). Across all cross-validation folds, the expert resistance surface consistently produced higher AIC and lower R^2^ values than the best-fitting empirical models, indicating a poorer fit to observed movement data. This demonstrates that the expert-derived resistance values did not fully capture the fine-scale behavioural constraints shaping blackbird movement in the urban matrix.

Under the repeated out-of-sample validation, the empirically parameterized model achieved modestly higher AUC and R^2^ values on average than the expert-based variants, suggesting that empirical calibration not only improves ecological interpretability but also enhances predictive capacity when evaluated under balanced data partitions. However, the difference in performance was only modest. Possibly, the smoother gradients of the expert maps may have generalized better to the data points included only in the test subset, while the empirically derived resistances, although more behaviourally realistic, were more sensitive to the specific spatial configuration of movement observations.

Taken together, these findings underline the complementary strengths of expert-based and empirically parameterized approaches. Expert maps provide stable, generalized estimates of urban permeability, while empirical calibration offers stronger process-based justification and quantifies where expert assumptions deviate from observed behaviour. Integrating both and using expert knowledge as an informed prior refined through empirical model selection can yield connectivity models that are both generalizable and ecologically grounded.

### 4.3 Limitations

Our modelling framework demonstrates that movement-proxy data can provide meaningful empirical information for calibrating urban connectivity models. The consistency of parameter estimates across cross-validation folds indicates that the parameterization procedure produces stable results even under varying subsets of data. This stability contrasts with the stronger fluctuations observed in validation metrics and suggests that parameter calibration may provide a more robust assessment of landscape permeability than single validation tests, which are more sensitive to the specific configuration of observation points. The ability to derive parameter uncertainty and to map areas of high and low model confidence further strengthens the transparency and interpretability of the approach.

Nevertheless, several limitations must be acknowledged. The use of movement-proxy data inherently introduces a degree of uncertainty regarding behavioural interpretation. Although the dataset captured active movements across the urban matrix, it may still integrate multiple behavioural states (e.g., commuting, foraging, or exploration), each associated with different spatial scales and sensitivities to landscape features. However, during the modelling, we strongly considered the spatial and behavioural properties inherent to our dataset and how to correctly reflect them in the connectivity model to counteract one of the main weaknesses outlined by Habrig and Fahrig (2025): the spatial mismatch between the model and the data used for validating it. Since we selected streets as sampling locations for the movement observations and the sampling season was before the juvenile dispersal, we were confident to assume that the vast majority of observed movements were directed movements between patches within the home range (McKellar et al. 2015; Dzialak et al. 2015).

A further limitation concerns the model’s sensitivity to spatial resolution. When the resistance surface was resampled at different grain sizes, the resistance estimates changed partially. This scale sensitivity is consistent with earlier findings that connectivity metrics depend on spatial resolution (Arponen et al. 2012), especially in highly urban areas (Lin et al. 2021). Identifying an ecologically meaningful resolution that corresponds to the perceptual range and movement scale of the focal species, therefore, remains a key requirement for robust urban connectivity modelling (Schooley and Wiens 2003). During the use of connectivity modelling and this parametrization procedure in particular, this could be addressed by performing explicit sensitivity analyses, systematically evaluating how changes in spatial resolution influence parameter estimates, model performance, and the resulting connectivity maps. Such analyses would help determine the optimal resolution for balancing ecological realism, computational efficiency, and transferability across spatial contexts.

The restricted number of parameter combinations tested is another limitation. While the factorial design of resistance and distance values was sufficient to detect major patterns, relevant intermediate combinations might not have been explored. A more continuous parameter search, or Bayesian optimization, could help identify finer-scale optima and improve model precision. Furthermore, the relatively coarse representation of some land-cover classes, particularly “streets,” likely contributed to the variability in resistance estimates. Incorporating finer-grained information such as traffic density, noise levels, or pedestrian intensity could yield a more realistic depiction of urban permeability (LaPoint et al. 2015).

Finally, our empirical calibration does not replace individual-based movement data such as GPS tracking or high-resolution telemetry, which remain the most direct means of quantifying animal movement and behaviour. However, such data are often logistically prohibitive or spatially limited. The strength of the presented approach lies in its ability to use simpler, opportunistically collected movement observations to infer realistic resistance patterns at a city-wide scale. By providing a transparent, statistically grounded framework for parameter estimation and uncertainty assessment, the method complements traditional expert-based and tracking-based approaches and can guide future data collection efforts toward critical areas of uncertainty.

### 4.4 Outlook

This study demonstrates how empirically derived movement parameters can enhance both the ecological realism and predictive ability of functional connectivity models in complex urban landscapes. By integrating direct movement observations into model calibration, we move beyond assumption-based parameterization and provide a transparent, data-driven framework that links landscape structure to observed movement behaviour. The resulting model not only performs better than expert-based approaches but also offers a stronger process-based foundation and a quantifiable measure of parameter uncertainty – an essential feature for applying connectivity models as decision-support tools in urban planning.

The methodological approach presented here is transferable to other species and cities, provided that movement-proxy data are available or can be collected opportunistically. The increasing availability of urban biodiversity observations through citizen science platforms, automated sensors, and camera-trap networks offers new opportunities to derive empirical resistance surfaces across taxa. Integrating these diverse data streams into statistical parameterization frameworks could lead to a new generation of empirically grounded, multi-species connectivity models that better capture the behavioural diversity of urban wildlife.

From an applied perspective, empirically parameterized connectivity models can inform the spatial prioritization of green infrastructure by identifying corridors and stepping stones that most effectively support functional movement. Integrating empirical calibration into the urban landscape planning workflow offers a path toward more evidence-based, ecologically meaningful connectivity assessments that align with international biodiversity and urban resilience targets.

## Notes

### Competing Interest Statement

The authors have declared no competing interest.

### Summary of Updates

This version presents a conceptually updated connectivity modelling framework. It includes new analyses that validate the model and demonstrate its practical value. The manuscript text has been substantially revised and streamlined to improve clarity and readability.

### Funding

- Bundesinstitut für Bau-, Stadt- und Raumforschung: Green Urban Labs II

