## Supplementary Information for "From expert opinion to empirical evidence: Data-driven parametrization of urban connectivity models using movement-proxy data"

### S1 Observational data on the common blackbird

**Dataset 1 – Blackbird presence/absence data for determination of used resource types**

Data on the presence and absence of the common blackbird was collected by performing a visual and acoustic survey along a gradient of greenness and traffic density in Munich in May and early June 2024. The aim was to gather information on the occurrence of blackbirds in Munich at a small spatial scale. To account for the variability of the urban landscape, we sampled across gradients of greenness along which the probability of observing a bird is expected to differ. Similarly, we aimed to account for the disturbance by cars and, therefore, selected points along a gradient of traffic density.

We used the land cover map described in Appendix 1 and data on car density from the navigation company TomTom (TomTom in press) to sample observation points. First, we derived the percentage of vegetation cover around all streets using a moving window with a radius of 300 m. Then, we created 3 equally sized classes of traffic density based on the measured car hits by TomTom and 3 equally sized classes of vegetation cover. From the combined 12 classes, an equal number of observation points was randomly sampled. This resulted in the selection of 104 observation points. A trained ecologist visited the sampling points on weekdays between 6:00 and 11:00 a.m. At every sampling point, the ecologist recorded the time and temperature. Then, for 10 minutes, the ecologist visually and acoustically monitored for the presence or absence of the common blackbird *Turdus merula* at the observation point.

**Dataset 2 – Blackbird movement data for parametrizing connectivity model**

Data on the movement of blackbirds was retrieved by observing blackbirds over sections of streets. The aim was to obtain information on the presence and absence of moving blackbirds throughout the urban landscape. To account for the variability of the urban landscape, we sampled gradients of greenness and tree presence along which the probability of movement probably differs. The distance to the city centre is a proxy of the intensity of urbanization and was therefore included as an additional gradient for sampling to account for the combined effects of dense urban areas on bird occurrence and movement.

The data was collected in June 2021. Trained ecologists recorded blackbirds on 217 street transects bordering the 103 squares described in Mühlbauer (2021). Thus, the movement covers the same gradients of square size, distance to the city centre and presence of trees. As blackbirds could be safely identified up to 60 m away, the streets were divided into transects of 10 x 60 m sections. If possible, three sections were sampled around each square. If the squares were smaller, the number of street sections was reduced to two or one. The ecologists observed the transects between 6 and 11 a.m. on weekdays and counted blackbirds for 4 minutes. The number of blackbirds flying over a street section summed up for every section. The time, date, number of pedestrians, wind speed, and temperature were collected as variables possibly influencing the number of birds flying. In the following, we use the presence/absence of movement derived from the abundance. As the movement of blackbirds was recorded over 60 x 10 m transects, we average variables across the area of the transect when trying to explain the presence and absence of blackbird movement over one transect.

### S2 Creation of the land cover map

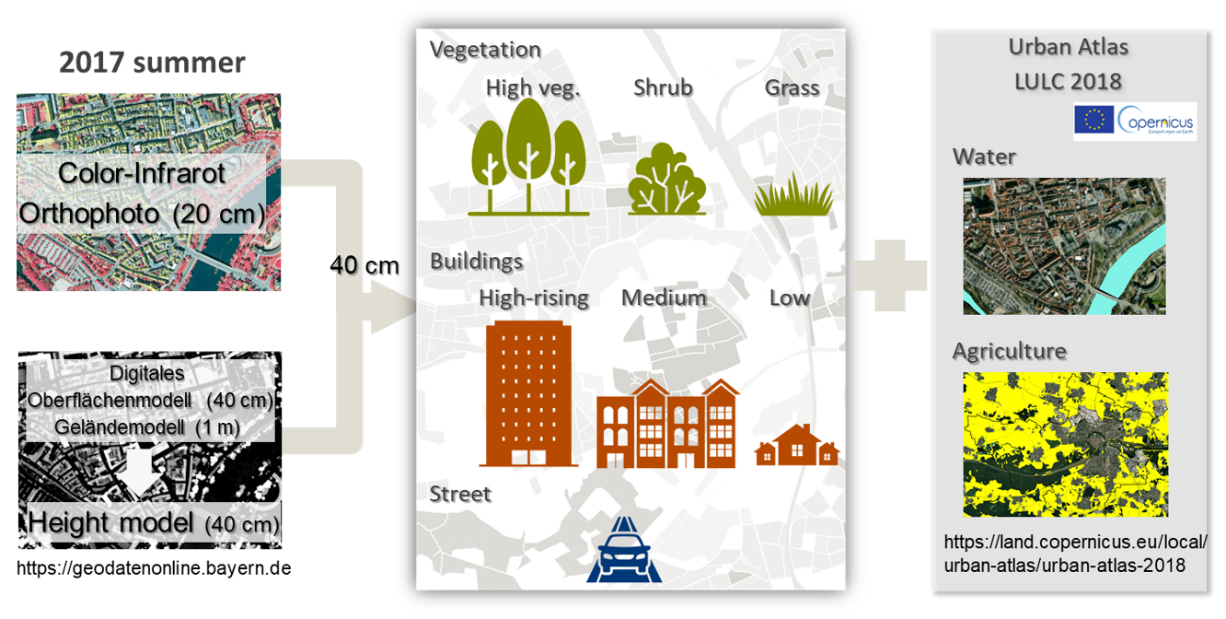

Figure S1 Overview of creating land-use land-cover map

We created a land-use land-cover map (LULC map) of Munich as input for the connectivity modelling. The LULC map should depict the barriers that animals face in the city and the resources and vegetation that attract them (Pundsack et al. 2025). Regarding vegetation, blackbirds use trees, shrubs, and grasses for nesting, foraging, and shelter (Snow 2008). However, they are used differently – especially grass surfaces mostly serve for collecting earthworms, whereas grass and shrubs provide shelter. Nevertheless, prior studies have found that trees especially impact bird movement (Tremblay and St. Clair 2009, 2011); therefore, we decided to not only differentiate between grass and other vegetation, but also between shrubs and trees.

Streets are known to represent barriers to the movement of birds, therefore, they were required as an additional land cover that probably acts as a barrier to blackbird movement (Tremblay and St. Clair 2009). For buildings, the relationship is not as well known, but prior studies suggest that buildings can act as a weak barrier to the movement of urban birds (Shimazaki et al. 2016). Additionally, knowing that blackbirds tend to fly quite low, we decided to include buildings and to build different classes of building height to account for barrier effects depending on the height of the building. Overall, these considerations required the acquisition of data that can differentiate vegetation from non-vegetated areas in the city and for a digital height model to account for the height of vegetation and buildings.

Digital surface and ground models (DSM and DGM) and orthophoto acquired in July 2017 for Munich were used for creating the LULC map. The most recent summer was selected for the orthophoto data. The spatial resolution is 40 cm for DSM, 1 m for DGM, and 20 cm for orthophoto. First, the normalized height model was created by subtracting DGM from DSM, resulting in a 40 cm resolution. The orthophoto was resampled to 40 cm resolution, and normalized difference vegetation index (NDVI) was created by red and near-infra-red bands. Second, vegetation pixels were divided from non-vegetation pixels by a threshold of 0.25 of NDVI. The threshold was manually decided while comparing the classified pixels with orthophoto. Third, by height, tree pixels were classified from vegetation pixels when normalized height was over 3 m, shrub pixels between 0.5 and 3 m, and grass pixels lower than 0.5 m. But, for the agriculture area, pixels between 1.2m and 3 m were selected for the shrub to exclude crops and pasturage taller than the grass of the residential area (i.e., 0.5 m). Beforehand, agricultural areas were selected from Urban Atlas LCLU 2018. Fourth, non-vegetation pixels were classified into high-rising buildings with a height over 18 m, medium-rising buildings between 10 and 18 m, low-rising buildings between 1.5 and 10 m, and streets lower than 1.5 m. For the non-vegetation pixels, Region Group and Nibble tools in ArcGIS Pro were applied to remove small pixel groups falsely classified: for example, tall cars on the street falsely classified into low-rising buildings. The Region Group identifies the connected pixels with the same class as individual groups, and the Nibble replaces the class of the selected area with the class of their nearest neighbor. Fifth, water was added to the classification after being selected from Urban Atlas LCLU 2018. Lastly, the classification quality was checked with random points for each class. The points were randomly created from the test sample polygons, which were manually selected using orthophoto.

Table S1 Classes of land-use land-cover map

| **Classification** | | **Height in**  **urban area** | **Height in agriculture area** | **Data source** |
| --- | --- | --- | --- | --- |
| Water | | - | | Urban Atlas LCLU 2018 |
| Agriculture | | - | 0 - 1.2 m | Urban Atlas + DSM/ DGM |
| Vegetation | Grassland | 0 - 0.5 m | - | Orthophoto  + DSM / DGM |
|  | Shrub | 0.5 - 3 m | 1.2 – 3 m |  |
|  | Tree | over 3 m | |  |
| Artificial surface | Street (or bare soil) | 0 – 1.5 m | |  |
|  | Low-rising building | 1.5 - 10m | |  |
|  | Medium-rising building | 10 - 18m | |  |
|  | High-rising building | over 18 m | |  |

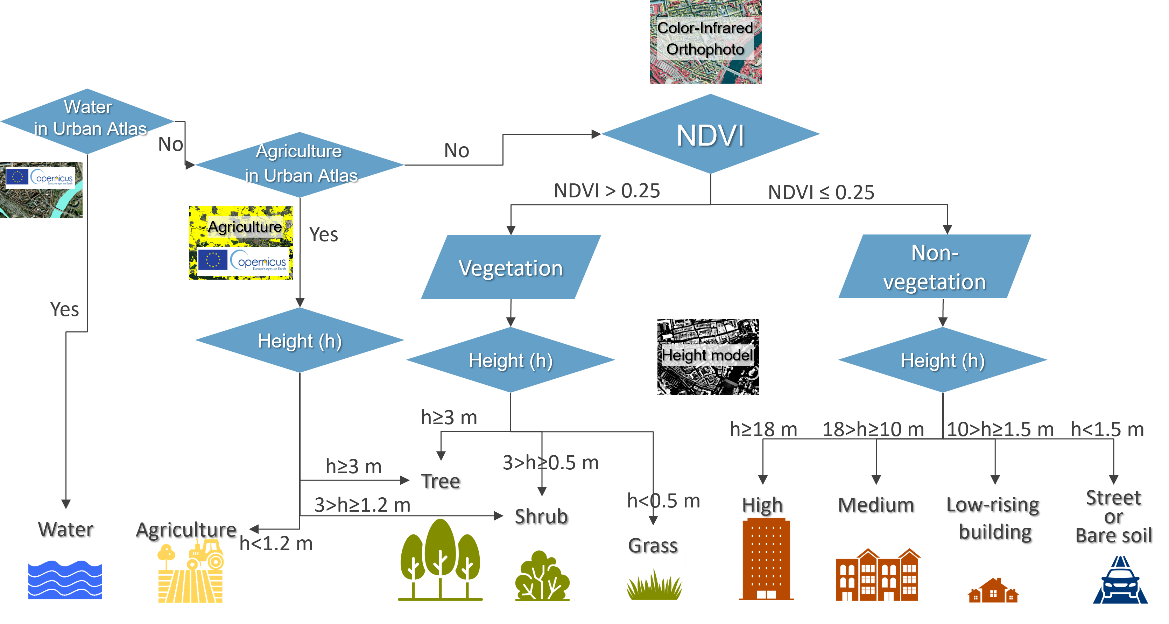

Figure S2 Flowchart of land-use land-cover mapping

Table S2 Evaluation of classification accuracy of Munich map

| **Class number** | **Class name** | **Sensitivity** | **Specificity** | **Balanced accuracy** |
| --- | --- | --- | --- | --- |
| 1 | Grass | 0.74 | 0.98 | 0.86 |
| 2 | Shrub | 0.72 | 0.97 | 0.85 |
| 3 | Tree | 0.80 | 0.96 | 0.88 |
| 4 | Street | 0.70 | 0.96 | 0.83 |
| 5-7 | Building | 0.99 | 0.76 | 0.88 |
| 8 | Water | 0.78 | 1.00 | 0.89 |
| 9 | Agriculture | 0.81 | 0.89 | 0.85 |
|  | **Mean** | **0.79** | **0.93** | **0.86** |

### S3 Detailed description and justification of parameter selection in case study

*Part 1: Resource map*

Step 1 aims to map the occurrence of resources that the animal could move to and that could form nodes in the subsequently built landscape graph. However, to statistically determine the resource types, we first had to identify which resource types could at all provide resources to the common blackbird, furthermore, the minimum home range radius of the common blackbird and a suitable modelling resolution.To select the input variables in *Step 1* to identify potential resource types, the minimum home range radius and a suitable modelling resolution, we followed these considerations:

***Potential resource types:*** Grasses, shrubs and trees are used by blackbirds for feeding and/or nesting (Snow 2008) and could be differentiated in the initial land use land cover map. Therefore, all three resource types are considered potential resources in further analyses.

***Minimum home range radius***: We used a 100 m radius for the home range to reflect blackbird foraging movements during the breeding season, when activity is concentrated around nest sites. While Ferry et al. (1981) report average male home ranges of up to 180 m in urban areas, analyses of the movement distances demonstrate that common blackbirds rarely ever move more than 80 m of their nest, especially during breeding season (Török and Ludvig 1988, Snow 2008). By selecting 100 m, we ensured that the home range radius reflects the most intensively used core area, which aligns with the assumption that the model captures frequent, local movement, not rare exploratory excursions.

***Modelling resolution:*** Considering the home range radius of 100 m and computational power, we aggregated the land cover map with an initial resolution of 40x40 cm at a 10 m resolution by extracting the percentage of each land cover class per 10 m raster cell. We selected a spatial resolution of 10 m to balance ecological realism and model tractability. A 10 m cell size captures fine-scale heterogeneity relevant to blackbird foraging decisions (e.g., presence of shrubs or small lawns) while aggregating sub-meter resolution data (0.4 m) to a scale more relevant to the perceptual and locomotor abilities of blackbirds. For the great tit, a song bird whose home range is substantially smaller than that of the common blackbird, it has been shown that the spatial occupation can be resolved up to a few meters (Baldan and Van Loon 2022). Therefore, we conclude that 10 m cells can account for the spatial scale at which the common blackbird perceives its environment and directs its movement.

Moreover, our 10 m resolution ensures that multiple resource cells can fit within a single home range radius (100 m). This resolution is consistent with or finer than that used in previous blackbird modelling studies (Hostetler and Holling 2000, Von Dem Bussche et al. 2008, Pellissier et al. 2012). We verified that key features such as buildings, roads, and vegetation remained ecologically distinguishable at this scale. We acknowledge that alternative resolutions could affect model outcomes and address this in Appendix 7 on sensitivity analyses.

***Statistical parametrization of resources***: Next, we aimed to identify the land covers that increase the probability of blackbird occurrence. For this, we used blackbird presence/absence observations in Dataset 1. However, we also wanted to identify areas where the minimum amount of resources required for the blackbird to establish is fulfilled. The reason for the latter goal is to exclude areas from connectivity modelling where the total amount of resources is too restricted for the blackbird to occur at all. The simplest approach for determining the minimum amount of resources necessary without having to threshold a continuous variable of habitat suitability is Classification and Regression Trees (CARTs). We decided not to employ more complex machine learning techniques like Random Forests, despite their ability to handle categorical response variables and demonstrate high predictive performance, due to the difficulty of interpreting the results (Cutler et al. 2007, Barnard et al. 2019). Additionally, we aimed to obtain simple classes for landscape description, where CARTs are easier to handle and provide clear and comprehensible results (Pesch et al. 2011, Mimet et al. 2019).

Furthermore, CARTs were chosen for their capability to handle non-normally distributed predictors, categorical response variables, as well as nonlinear and complex relationships between predictors and responses, including interaction effects, as evidenced in previous studies (De’ath and Fabricius 2000, Bel et al. 2009), making them a robust choice for our analysis (Pesch et al. 2011, Mimet et al. 2019).

When creating the CARTs, we implemented a pre-pruning criterion to prevent overfitting of the tree model by defining a minimum of eight observations per terminal node. Pre-pruning constrains tree growth by setting a threshold for observations in each terminal node, ensuring the model remains generalizable. During tree construction, potential splits at each node are evaluated, but a split is only made if each resulting terminal node can contain the specified minimum number of observations. If not, the node remains terminal, thus avoiding the creation of overly specific nodes that might capture noise rather than meaningful patterns. This technique regularizes the model and ensures that it focuses on substantial relationships by reducing the risk of overfitting on small data fluctuations. The threshold of eight observations per leaf was chosen based on previous studies (Debeljak et al. 2001, Pesch et al. 2011), the dataset size, and visual assessments.

Using the CARTs, we modelled observed presence and absence of the common blackbird in Dataset 1 as a function of the proportion of trees, grass and shrubs in a 100 m circle around each observation point. Presence (1) and absence (0) were provided as classes. CARTs were modelled using the rpart package (Therneau and Atkinson 2022) in R. The CART model then determined the amount of all three resource types that is required in a 100 m circle for the presence of the common blackbird.

After developing the CART model, we evaluated its performance using a confusion matrix, which summarizes correct and incorrect predictions across classes (presence/absence of the species). From this matrix, we calculated accuracy, sensitivity, specificity, and kappa, each capturing different aspects of predictive performance. Sensitivity assesses the model’s effectiveness in identifying positive cases, while specificity measures its ability to avoid false positives. Kappa indicates agreement between observed and predicted classifications while adjusting for random chance, making it more reliable than accuracy alone, particularly with imbalanced data.

When applying the CART model to the total study area, the city of Munich, we identified the areas where the amount of resources was higher than the minimum amount of resources necessary for the occurrence of the common blackbird determined by the CARTs. The aim of this step was to identify the areas where the resource amount was too low for the blackbird to occur at all, and that could therefore be excluded from connectivity modelling. However, we could not really identify any areas, where, within a radius of 100 m, the modelled minimum requirements of resource amount were not met. Thus, we decided to model connectivity to all areas of Munich.

***Building the resource map:*** The resource map provides information on which areas could attract the target species. Thus, we used the information on which resource types positively contributed to the occurrence of the common blackbird according to the CARTs. However, we had information on the proportion of the different resource types per 10 m cell. Thus, we had to set a threshold to decide how much cover of a resource types makes a cell attractive for the common blackbird.

We assumed that bird use resources patches of small size. However, there is a trade-off between including all resources possibly attractive and computational power required to run the cell-based connectivity model. After visual inspection and performance tests, we decided to set the threshold at 20m², which is not too small considering that this is the size of a well-grown tree (Konijnendijk 2023). All pixels containing more than 20m² of the resources were mapped in the resource maps and designated as nodes in the connectivity analysis.

*Part 2: Resistance values and node isolation distance*

After determining where the resources are located, we calculate the connectivity between those resources. However, this requires us to determine the resistance values for different urban land cover types as well as the typical distance that the blackbird moves without using stepping stones, i.e. the node isolation distance. To determine the resistance values and the node isolation distance, we first created resistance and node isolation distance scenarios, then modelled the connectivity for each combination of scenarios and then determined the resistance values and node isolation distances that best explained the observed movement.

***Resistance and node isolation scenarios:*** Resistance values in connectivity models summarize several behavioural decisions, including movement preferences, avoidance behaviour, energetic consideration and perceptual limitations, and mortality (Zeller et al. 2012). These values are rarely measured directly and are typically derived through expert knowledge (Zeller et al. 2012, Riordan‐Short et al. 2023). Direct estimation from movement data is only possible when using fine-scale trajectory models such as step or path selection functions (SSFs), which require detailed tracking data (Zeller et al. 2016). The aim of our parametrization procedure is to leverage scenario building and model selection to derive resistance values (Verbeylen et al. 2003). Although resistance was not directly derived from movement speed or directional persistence, our model selection approach ensures that the resistance scenarios best explaining observed movement patterns are retained. However, defining all ecologically possible resistance scenarios is an important prerequisite for deriving resistance values of high predictive power.

To assess the barrier effect of urban elements on blackbird movement, we created different *resistance scenarios* in which we assigned resistance values to different land cover types based on their possible impact on bird movement. In all scenarios, resources were assigned a resistance value of 1. We lacked observation points close to water and agriculture and, therefore, could not robustly test their possible resistance values. We therefore attributed them a resistance value of 10 due to their open nature, which offers a certain resistance to movement (Tremblay and St. Clair 2009). Similarly, vegetation not identified as a resource was assigned a resistance value of 10.

We based resistance values of anthropogenic structures (e.g., buildings, roads) on their known barrier effect to songbird movement and assigned them a possible range of higher values (up to 1000) to reflect steep energetic and predation risks associated with crossing impervious or elevated surfaces (Tremblay and St. Clair 2009, 2011, Shimazaki et al. 2016, Ersoy et al. 2019). We decided to include *resistance scenarios* with values up to 1000 to include very strong barrier effects, especially of streets and buildings. Nevertheless, in certain scenarios, the maximum resistance value is 10 or 100, leading to *resistance scenarios* with a weaker barrier effect of buildings and streets. We ensured that the resistance of taller buildings was not lower than that of shorter ones, although we did not establish prior assumptions regarding the relative resistance of buildings compared to streets. This approach resulted in 24 different *resistance scenarios* for testing that include all combinations given the restriction that the resistance of lower buildings cannot be higher than that of higher buildings and the defined range of values (Figure S2). For each *resistance scenario*, we created a resistance map from the 10 m resolution land cover map. Each map was computed by summing the weighted resistance values of land cover types present in each cell of the 10 m land cover map.

We established four node isolation distances – 30, 60, 90, and 150 m – based on literature concerning blackbird movement and home range sizes (Ferry et al. 1981, Török and Ludvig 1988, Cresswell 1999).

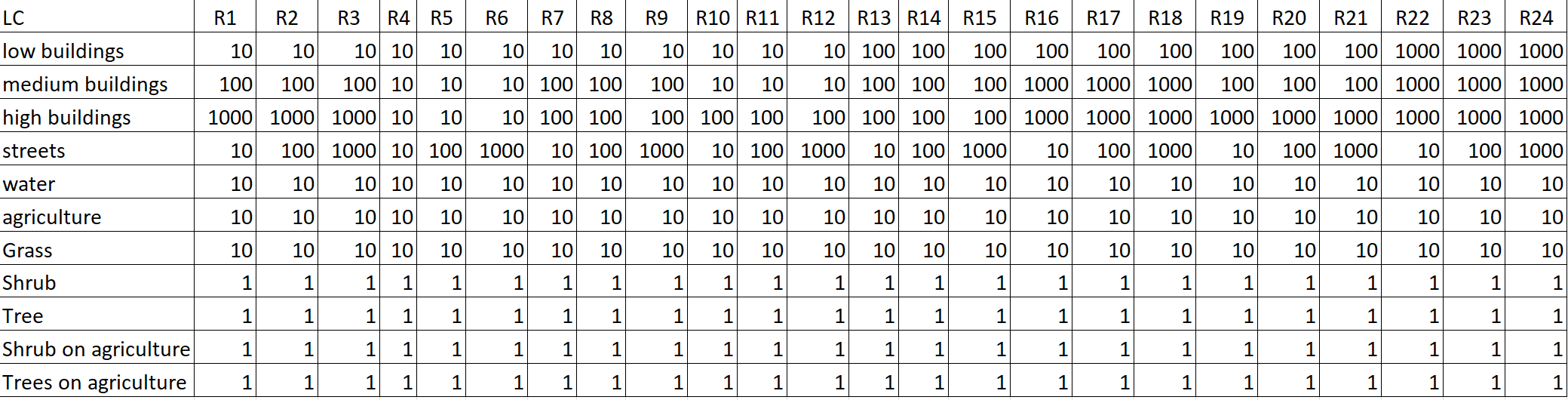

Table S3: Resistance scenarios to be tested in the case study

***Graph modelling:*** To build the graph-based connectivity model, we used the R package graph4lg (v. 1.8) (Savary et al. 2023) that utilizes the program Graphab (v. 2.8) (Foltête et al. 2012). The nodes of the graphs were all the cells containing at least 20 m^2^ of the resource types determined in *Step 1*. For each of the 24 resistance maps, we then created a set of links with a planar topology. For each of the 24 link sets, we created four graphs that were thresholded by the cost-converted value of the four maximum patch isolation distances (30, 60, 90, and 150 m) (details on set parameters in Table S4). The choice of the parameter ensured that we calculated link-based Betweenness Centrality on a cell-based landscape graph (Figure S3).

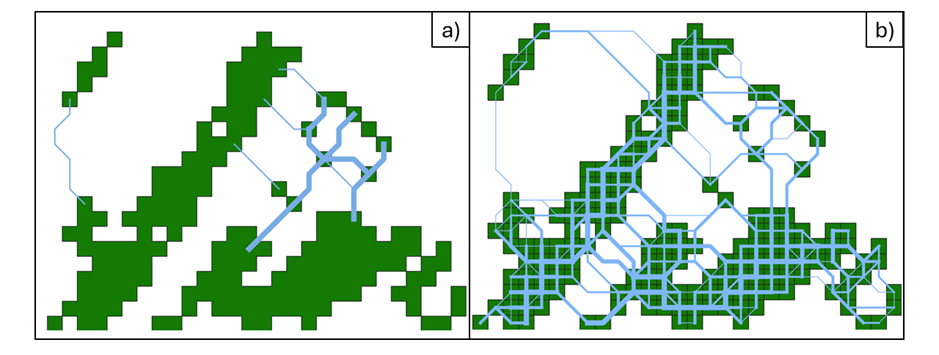

Figure S3: Figure 1: Patch-based (a) and cell-based connectivity (b). In the patch-based landscape graph (a), the green resource patches form the nodes. Two resource patches are connected by least-cost paths (blue) (a). In the cell-based landscape graph (b), the green resource cells form the nodes (b). Two resource cells are connected by least-cost paths (blue). The line width of least-cost paths in a) and b) reflects the Betweenness Centrality of the links

Table S4: Parameters set to model the landscape graph. The functions refer to the graph4lg R-package (Savary et al. 2023) and to the command line functions of Graphab (Clauzel et al. in press). command line functions are indicated by --function

| Function | Parameter | Justification |
| --- | --- | --- |
| graphab_project() | maxsize = 10 | We wanted to derive a cell-based connectivity model and, hence, set the maximum size of each node to the resolution of the input map. |
| graphab_link() | topo = planar | We worked with planar graphs to reduce computation and to account for the fact that most animals use stepping stones to movement. |
| graphab_graph() | thr = node isolation distance | We pruned the linkset by the 4 node isolation distances set before, resulting in 4 graphs for each node isolation distance. |
| graphab_graph() | cost_conv = TRUE | We pruned the linkset by the cost-converted node isolation distance to omit links that pass areas of extremely high resistance for which we assume a very low probability of movement. |
| graphab_metric() | metric = BC | We aimed to calculate the Betweenness Centrality. When deriving the Betweenness Centrality, it is calculated for both the nodes and the links. |
| graphab_metric() | dist = node isolation distance | To ensure consistency within the connectivity model, we used the node isolation distances by which the graphs were pruned before. |
| graphab_metric() | *α* = 0.01 | we assumed that those distances represent the maximum patch isolation distance. |
| graphab_metric() | a = size of node | No further data was available on the capacity of our nodes and the assumption that capacity scales with area seemed robust, therefore, we defined the capacity *a* as the patch area. |
| graphab_metric() | β = 1 | For the choice of the β-parameter in the metrics calculations that determines the weight between the distance-term and the patch-capacity term we did not want to put too much weight on the patch capacity and therefore set it to 1. |
| graphab_metric() | cost_conv = TRUE | We again used the cost-converted node isolation distances to ensure consistency within the connectivity model. |
| --interpol | res_map = 10 | We wanted to derive an interpolated connectivity map at the initial modelling resolution. |
| --interpol | d = node isolation distance | To ensure consistency within the connectivity model, we used the node isolation distances by which the graphs were pruned before. |
| --inteprol | p = 0.01 | To ensure consistency within the connectivity model, we used the probability that was used when calculating the metric before. |
| --interpol | multi_d = patch isolation distance + 20 | For the interpolation, we included the BC-values of all links within the given node isolation distance plus 20 m to include all patches that could have a remote impact on the local connectivity. |

***Interpolation of Betweenness Centrality:*** We interpolated the Betweenness Centrality (BC) of the links using the metric interpolation tool of Graphab v. 2.8.6 (Clauzel et al. in press). However, the default use of the tool only allows interpolation of metrics that are calculated for the nodes – the interpolation from the links is not available in the standard functions. Therefore, we introduced additional steps to be able to perform this interpolation.

First, we calculated the BC of the links using the graphab_metric-function in the graph4lg R-package (Savary et al. 2023). Then we assigned the BC-values to the linkset-shapefile that contains the least-cost paths between the nodes. We rasterized the linkset imprinting the respective BC-values using the rasterize-function in the terra-package (Hijmans 2023) to a 10 x 10 m raster-layer. Within the rasterize-function, we summed up the BC-values of all the links that passed through the same raster cell. Thereby, we derived a raster-layer showing the sum of the BC-values of all the links touching a raster cell, whereas cells without link remained empty.

To interpolate this BC-raster-layer, we created a new Graphab-project only meant for interpolation. The nodes were set on all the raster cells that had a BC-value. Therefore, the interpolation graph had more nodes than in the initial graph for which the BC-value was initially calculated. Using the same resistance map as for the initial graph, we calculated the linkset and pruned it by the same node isolation distance as before. Then, using again the graphab_metric-function from graph4lg, we calculated the BC-value of all nodes in this interpolation graph. In the attribute table of the interpolation-graph, we replaced all the newly calculated BC-values by the values extracted from the rasterized BC-raster-layer. Finally, we used the interpolation-function from command line to interpolate the BC-values from the nodes of this manipulated graph. Overall, this allowed us to derive a raster map with the BC-values of the links being interpolated for the entire study area.

### S4 Testing parametrization results across size of dataset

*Methodology*

If many observations are required to determine the model parameters, this is can limit the applicability of the parametrization framework. Different technical approaches might be limited in the number of observations that could be required. For instance, the number of suitable locations to place a camera trap (LaPoint et al. 2013) could be limited or the number of observed roadkills relatively low (Koen et al. 2014). Thus, an estimate of how many observations are required to robustly parametrize the connectivity model can support researchers in estimating the sampling effort that has to be made derive sufficient datapoints.

Dataset 2 consists of 215 presence/absence observations of blackbird movement throughout the city of Munich. We could not increase the number of available observations, thus, we decided to take proportions of the datasets and determine the model parameters to compare them with the parametrization results obtained from the full dataset. Out of the full dataset, we performed a randomized stratified sampling considering the distance of the observation point to the city centre and the amount of surrounding green. We created one subsample consisting of 66 % of the observations out of the full dataset and one containing 33 % of the observations. Then, we performed the parametrization procedure as described in the main document using the presence and absence of the common blackbirds as a response variable in the 33 % and 66 % subsamples.

*Results*

The parametrization results for the resistance values were quite similar between the three datasets (Figure S3a). Especially when considering the full dataset and a 66 % random sample of this full dataset, there is nearly no difference detectable in the selected resistance values. For the node isolation distance, the results between the full dataset and its 66 % subsample are very similar. However, when using only 33 % of the observations, the selected node isolation distance is different from the full and the 66 % subsample (Figure S9b).

*Conclusion*

We found that the number of observations used to perform the parametrization procedure can impact the final parameter values. When working with 142 observations instead of 215, the parametrization results were very similar. However, especially regarding the node isolation distance, a reduction in the number of observations to only 71 led to different results.

We performed connectivity modelling across the city of Munich which constitutes a total area of 310 km^2^. Thus, 142 observations over the total area represent about one observations per 2 km^2^ when sampling effort is expressed as a spatial density. However, the number of observations required does not only depend on the total area considered. It also is strongly affected by the number of land cover types for which resistance values should be derived and the number of resistance scenarios because they ultimately impact the number of statistical models from which the best one is selected based on the observational data. Moreover, the resistance values can only be estimated for land cover types that are spatially close to many observations points. Otherwise, no impact of their resistance value on the connectivity values around the observations points can be detectable. Thus, the more land cover types are considered in the connectivity model, the higher the number of observations need to be.

We conclude that the selection of the resistance values is relatively robust to rather low sample sizes, whereas the estimation of the node isolation distance is rather sensitive to the number of observations used for parametrization. Therefore, we recommend considering literature values on typical movement distances that the target species performs between stepping stones when the number of available observations is low. Still, also under a restricted number of observations, the parametrization of the resistance values can be performed reliably.

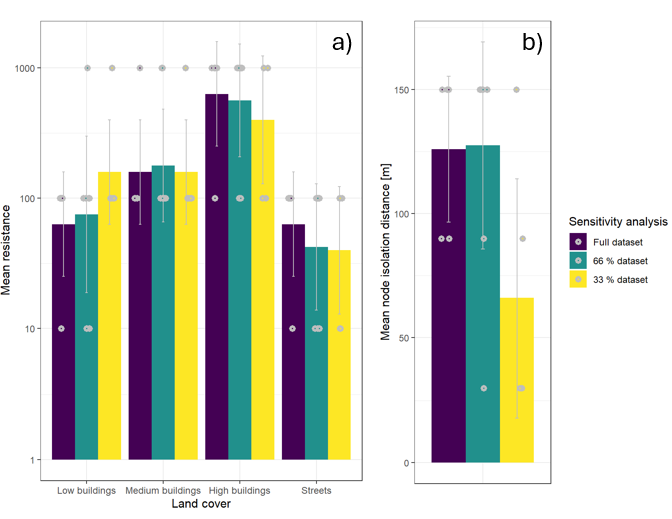

Figure S4: Parametrization results for urban land covers (a) and node isolation distance (b) when performing the parametrization with the full dataset (215 observations), 66 % of the dataset (142 randomly sampled observations) or 33 % of the dataset (71 randomly sampled observations)

### S5 Results of CART model

Table S5: Ability of CART to correctly predict observed presence and absence of blackbird and variable importance (VI) for the three predictors included (percentage of trees/shrubs/grass 100 m around the observation points).

| **Accuracy** | **Kappa** | **Sensitivity** | **Specificity** | **VI Tree** | **VI Shrub** | **VI Grass** |
| --- | --- | --- | --- | --- | --- | --- |
| 0.82 | 0.57 | 0.96 | 0.57 | 14.61 | 14.48 | 11.43 |

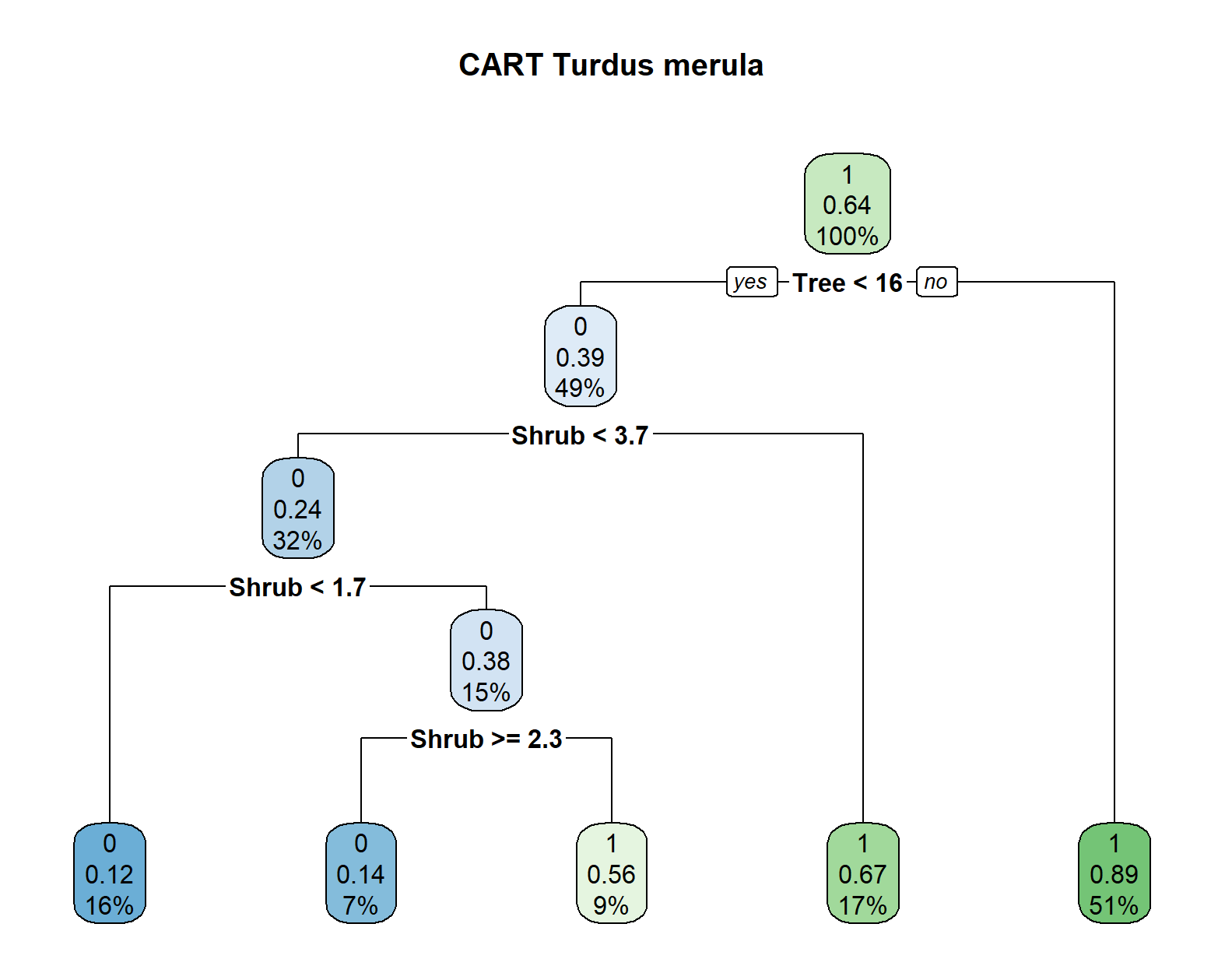

Figure S5: Classification and Regression Tree for classifying the presence (1) and absence (0) of blackbirds as a function of the coverage of shrubs, trees and grass around the observation points.

### S6 Sensitivity analyses

Sensitivity analyses can help identify the variables where a small change leads to a strong difference in modelling results (Cariboni et al. 2007, Rayfield et al. 2010). Moreover, they can support researchers in identifying for which variables a high effort should be made in their determination or in collecting field data (Cariboni et al. 2007). For example, if a variable is very sensitive to the value taken, it is worth spending more time on correctly identifying this variable compared to a variable that is relatively insensitive in the model. We performed several sensitivity analyses to test how the model results react to different assumptions and the number of observations.

First, we tested whether the minimum home range radius used to determine the critical resources in *Step 1* impacts the selection of resources applying CARTs. Then, we assessed how the spatial resolution, i.e. the size of the raster cells of the input map impacted the parametrization results for the node isolation distance and the resistance values. Furthermore, we tested how well different spatial resolutions performed when validating the results with an additional dataset.

**Sensitivity analyses on minimum home range radius**

*Methodology*

The minimum home range radius of a species is an important parameter that is fixed during the parametrization of *Step 1.* It is mainly used to assess in which radius around an observation point the amount of resources is extracted to predict the occurrence of the target species. It furthermore informs the *node isolation distance scenarios* tested during the parametrization of *Step 2.* However, since several values of the *node isolation distance* are tested during the parametrization of *Step 2,* we decided to only test how several values impact the selection of resources relevant for the occurrence of the common blackbird in *Step 1.*

We initially worked with a minimum home range radius of 100 m (see Appendix 3 for more details). However, we also tested minimum home range radii of 80 and 120 m in a sensitivity analysis. These values fit the movement ranges observed by Török and Ludvig (1988).

To perform the sensitivity analysis, we extracted the proportion of the three potential resources, grass, shrubs and trees, in the three radii (80 m, 100 m, 120 m) around the observations of the common blackbird denoted in Dataset 1. This was performed using the terra package and the function terra::extract. For each radius, we built one classification tree where the classes observed presence (1) and absence (0) of the common blackbird in Dataset 1 were modelled as a function of the three predictors, proportion of grass, proportion of shrubs and proportion of trees. We followed the pruning strategy as described in Appendix 3. For each radius, we furthermore assessed the performance of the respective classification tree by applying the fitted model to the training dataset. We created a confusion matrix and calculated the Kappa-value, accuracy, sensitivity, and specificity.

To assess how robust the outcome of the test was to several minimum home range radii, we compared whether the selected resources in the classification tree differed (see Appendix 4 for results of CART for 100 m radius). Additionally, we compared whether the minimum amount of resources required for the occurrence of the blackbird differed substantially when using different radii to create the tree. Finally, we compared the performance metrics Kappa, accuracy, sensitivity and specificity for the three values.

*Results*

The three classification trees are quite similar. The first split is always based on the minimum amount of trees around the observation point, the second split determines the minimum amount of shrub required for the presence of the common blackbird. This is then followed by smaller splits that differ partially between the three radii. Also the exact proportion of trees and shrubs required for the occurrence of the common blackbird differ between minimum home range radii – e.g. for 80 m minimum home range radius the requirements are either at least 17 % of trees or 1.9 % shrub coverage around the observation point. For the classification created for 120 m minimum home range radius, the minimum requirements are at lest 13 % tree coverage or a minimum of 2.9 % shrub coverage. In none of the final pruned trees, the proportion of grass occurs.

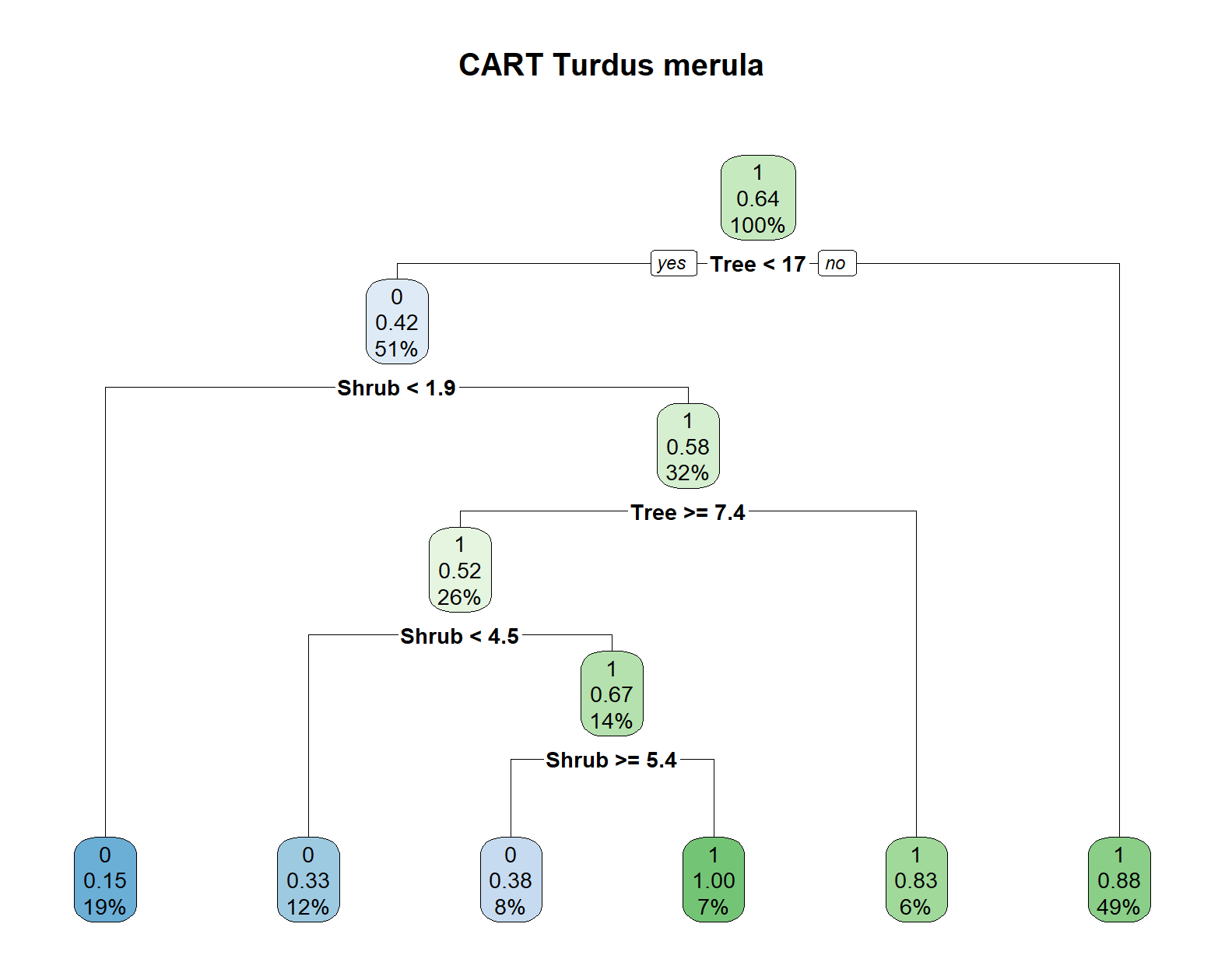

Figure S6: Classification tree for minimum home range radius of 80 m. This is part of the sensitivity analysis to test the value taken of 100 m

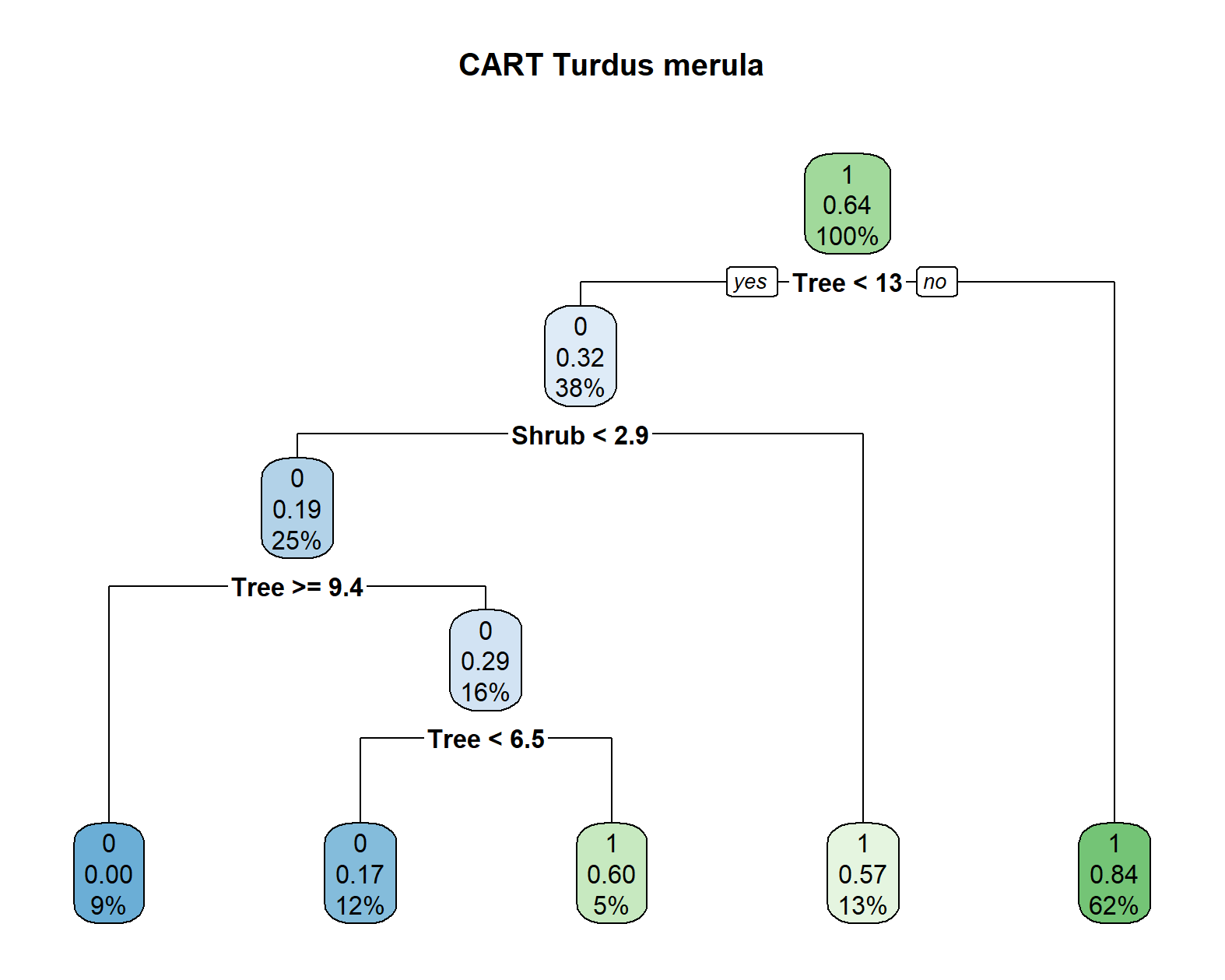

Figure S7: Classification tree for minimum home range radius of 120 m. This is part of the sensitivity analysis to test the value taken of 100 m

Table S6: Statistical performance indicators for classification trees using the three tested minimum home range radii (80 m, 100 m, 120 m). VIP indicate variable importance scores

| **Radius [m]** | **Accuracy** | **Kappa** | **Sensitivity** | **Specificity** | **VIP_Tree** | **VIP_Shrub** | **VIP_Grass** |
| --- | --- | --- | --- | --- | --- | --- | --- |
| 80 | 0.84 | 0.65 | 0.85 | 0.65 | 16.22 | 15.81 | 11.27 |
| 100 | 0.82 | 0.57 | 0.96 | 0.57 | 14.61 | 14.48 | 11.43 |
| 120 | 0.81 | 0.54 | 0.97 | 0.54 | 16.15 | 10.53 | 9.97 |

The performance metrics for the three radii are relatively similar with accuracy values ranging from 0.81 to 0.84 and Kappa-values ranging from 0.54 to 0.65 all indicating a good model fit beyond random chance. For the 80 m radius, the specificity is higher – this classification tree was better at predicting absences. The CARTs created from 100 and 120 m minimum home range radius performed better at predicting presences indicated by ther very high sensitivity values.

*Conclusion*

From the sensitivity analysis, we deduce that while the classification trees show certain differences when varying the minimum home range size, the overall results remain robust. Thus, similar resource types are shown to be relevant for the occurrence of the common blackbird and relatively similar minimum amount of resource required for the presence of the target species are identified. The statistical performance indicators suggest that the CART created from 80 minimum home range radius performed a bit better than those from the other radii – however, this difference was relatively small. Finally, since all CARTs predicted nearly the entire area of Munich to generally provide sufficient resources for the occurrence of the common blackbird, the exact minimum requirements were not used in the model. We only extracted the information that the common blackbird is positively associated to the amount of shrubs and trees. Since this information was constant across all tested minimum home range radii, we consider the choice of a minimum home range radius of 100 m a robust and well-grounded decision.

**Sensitivity analysis on cell size**

*Methodology*

The resolution at which modelling is performed is an important parameter that is determined during the parametrization of *Step 1.* It is probably also one of the most delicate methodological choices that has to be made during the modelling process. The cells should have a size such that one cell represents one location where within the cell only movements typical for encamped behavioural states are observed. However, to move between cells, the animal should need to perform the long, goal-directed movement that are modelled using the landscape graphs. Moreover, the cell size should be small enough to account for the spatial variation of the landscape. However, the spatial resolution used for modelling also strongly impacts the computational requirements. Also, depending on the study area, land cover maps of a spatial resolution below 10 m might also be difficult to be acquired (Pundsack et al. 2025). Hence, a very fine resolution suitable is both ecologically and computationally not suitable, whereas a coarse resolution will facilitate computations but can lead to overlooking ecologically relevant movement. As long as no very fine resolved tracking data is available to determine a suitable cell size, the cell size needs to be determined from knowledge on the species, its behaviour and its body size. Here, a sensitivity analysis can support researchers in assessing how robust the results are to the choice of this very important parameter.

For the main study, we performed modelling at 10 m cell size. However, we also tested model performance for 6 and 20 m spatial resolution. Since our initial land cover map had a resolution of 40 cm (Appendix 2), we had to aggregate to a cell size that is a multiple of 40 cm. Therefore, we selected 20 and 6 m cell size as reasonable resolutions above and below our initial choice of 10 m.

To perform the modelling for 6 and 20 m, we followed the same modelling steps as described in the main manuscript but we did not work with the cross-validation. We used the full Dataset 2 for parametrization and worked with a map of the respective resolution. We did not test the impact of cell size on the CARTs used to identify the resources in *Step 1* because the cell size does not impact the amount of resources extracted around an observation point. However, we tested the parametrization results obtained for the node isolation distance and the resistance values in *Step 2*. We considered the model to be robust to the cell size if similar parametrization results were obtained across the three cell sizes (6 m, 10 m, 20 m).

*Results*

When parametrizing the connectivity model at different spatial resolutions, the parametrization results for both the resistance values and the node isolation distances vary for the different cell sizes (Figure S7). Most importantly, the relative order of resistance values shifts between spatial resolutions. For 20 m resolution, the resistance to streets is highest, whereas the resistance to low and medium height buildings is lowest. Highest resistance of high buildings is detected at 10 m resolution, whereas the resolution of low buildings and streets is similarly low. When modelling with a cell size of 6 m, streets show the lowest resistance, whereas both, medium and high buildings show a high resistance (Figure S7). For 20 and 6 m modelling resolution, relatively low node isolation distances are detected, whereas for 10 m resolution, it is substantiall higher (Figure S7).

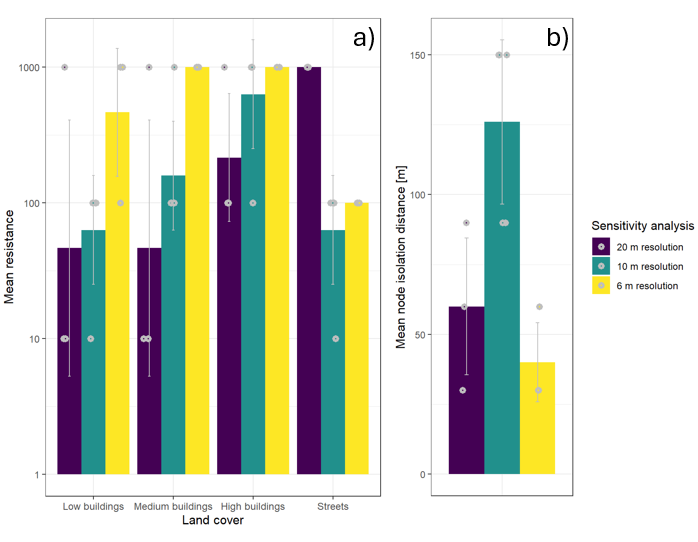

Figure S8: Resistance values for urban land covers (a) and node isolation distance in m (b) for the three cell sizes tested in the sensitivity analysis (20 m, 10 m, 6 m)

Importantly, the significant effect of connectivity on the observation of moving blackbirds can only be detected when modelling at 10 m cell size (Table S11).

*Conclusion*

Comparing the parametrization and modelling outcomes using different spatial resolutions demonstrates that both parametrization results and the resulting connectivity model are sensitive to the chosen modelling resolution. This is important to consider and points to the need to make a well-informed choice of the modelling resolution.

However, our results also demonstrate that the chosen modelling resolution of 10 m is the one where the effects of connectivity on blackbird flights are most clearly detectable. At 6 m resolution, the effects of resource availability are more detectable, but the lower AIC and R^2^ values clearly demonstrate that this resolution and the parametrization results are less suitable for predicting blackbird movement. When performing the connectivity modelling at 20 m, the overall performance metrics are comparable to those of 10 m cell size. However, the individual effects demonstrate that this resolution does not well capture the local effects of connectivity. Possibly, this increased performance is due to considering resource availability at a larger spatial scale because it is assessed from larger cell sizes. Thus, this can take up important parts of the variability explaining connectivity.

Generally, we conclude that the modelling resolution is a very important parameter that can alter the modelling and parametrization results and should therefore be carefully chosen. We recommend to perform sensitivity analyses and do a thorough literature review to determine a suitable modelling resolution. Finally, the results of our sensitivity analysis support the selection of a 10 m cell size in our model.
